# An ancient evolutionary calculus for attention signaling retained in modern music

**DOI:** 10.1101/2025.09.28.679029

**Authors:** Gregory A. Babbitt, Ernest P. Fokoue

## Abstract

What is in a song? Although music has long been subjected to philosophical and scientific study, the mechanism(s) of fitness or quality signaled through music has not. As music and dance functionally aim to hold an observer’s attention, we investigate the features of sound that stimulate attentive behavior in an audience. We propose a calculus of attention to characterize a singer’s *f*(*x*): **C**ontrol of vocal performance, *f*(*x*); **E**nergy within vocal performance, *f′*(*x*); and **S**urprise or change in direction of performance, *f′′*(*x*) (CES). We develop theory and software to, (A) extract CES, (B) dynamically map trajectories within the ternary space of CES, and (C) measure song stability (i.e. quality) and individuality. We validate that both human music and animal vocalization maintain high stability in the space of CES, with professional musicians being most stable. CES measurement also reflects differences in expert vs. novice singing ability in both humans and canaries as well as constraints during live audience interaction as well. We conclude that quality signaling via this calculus of attention reflects a cognitive focus upon motion in the environment created by selective pressures of intra/interspecific conflicts formed after the Ediacaran-Cambrian boundary and still retained in music today..

**Lay Summary:** Most animals (including humans) pay attention to specific audiovisual stimuli within their environment. Here we explore common features of these stimuli and propose a single mathematical framework (with software) for detecting and measuring the quality of performance regarding attention signaling in the context of sound. We show that musical sounds are exceptionally attractive to our attention and that attention signaling in music probably has ancient evolutionary roots that connect to when early life forms started to evolve swift directed movement and the sensory capacities to detect it.

## 1 Introduction

What is ‘song’? And what is contained within a ‘song’? Why have we applied the word ‘song’ not only to our own human vocal-social behavior, but also to the vocal behavior of some other animal groups as well? Why are vocalizations in animal groups such as sub-oscine passerines (i.e. songbirds), cetaceans (i.e. humpback whales), some primates such as Hylobates (i.e. gibbons and siamangs), and some anurans (i.e. chorus frogs) viewed as more musical to our ears than the vocalizations of most other animal groups? Within songbirds, why are some species such as Nightingales, Song Sparrows, and Lyrebirds considered especially musical? Is this difference purely an artifact of human perception or is there a tangible acoustic difference that we can detect with a technological approach? If singing is the oldest form of human musical behavior, then what can a mathematical comparison of human singing to other kinds of animal ‘song’ inform us about what music actually is and perhaps what it is not?

The origin, purpose, and nature of music have long been viewed through very different intellectual perspectives. Natural philosophers have repeatedly observed that music has potential roots in both cosmic and Earthly forms of nature [42][93], and like mathematics[94], music also seems to possess elements of both an internally-represented language in our minds as well as an external physical reality that we can directly and objectively measure. More recently scientists across diverse fields have also explored the origin and function of music. Consensus across scientific disciplines might conclude that music contains features very close to and yet predating human language [30][29][64][72], is culturally diverse and yet rooted to some degree in genetics [10][11][38][60], has structural features (e.g. isochrony, improvisational dueting, harmonic overtone series, intentional composition, universal tempo) that occur in some animal vocalizations outside humans [7][17][22][23][18][6][34][35][69][73][80] and can even can occur spontaneously within artificial intelligence [43]. Music fosters human social connection in the absence of shared language [70][76][81][95], and evokes stimulation and development across large parts of the human brain [3][9][31][37][75][86][92]. Anthropologists and biomusicologists have linked the evolution of music to Darwinian fitness [2][36][61] and the rise of bipedalism [81], the formation of prosocial behavior in human societies [65][76] and perhaps molecular systems of interaction as well [5][4]. Musical behavior has also been linked to the evolution of self-awareness and the ability for cognitive abstraction via theory of the mind [23][81]. One perspective that has been less often applied to the subject of music is that of behavioral ecology. Here, we would begin with basic questions about what fitness advantage is ultimately conferred to individuals engaging in musical behavior [36], and more specifically, what are the exact proximate mechanisms that signal individual quality (i.e physical fitness) in the performance display of musical behavior. While scientists have proposed many plausible ultimate evolutionary biological and cultural drivers of the evolution of musical behavior, to our knowledge, the proximate mechanisms by which individual quality is honestly signaled through musical performance has not yet been thoroughly investigated.

Whether or not a song is originated by a human voice, at its most basic level, a song (and dance) most often functions to hold the attention of one or more observers [47]. Therefore, we might define the proximate function of song to simply hold the attention of others. In this respect, it shares a common function with all other forms of behavioral display. Attention or focus in the brain is controlled largely by the dorsal and ventral attention networks, an evolutionary ancient but recently expanded set of connected brain regions [83][91][90][62] with strong anti-correlated activity with the mammalian default mode network [46][27], a set of connected brain regions activated during resting states and responsible for internal mental representations of self-awareness [54]. Interestingly, while music has features that stimulate attention [49] and emotion [96], it also has been shown to stimulate the default mode network as well [85], perhaps suggesting some interesting but as yet unexplored causal links between the evolution of human music and the several largely unexplained brain size expansion events in hominid evolution [28] [78].

When we consider musical performance within the context of other types of animal behavioral display, we can see that there are some potential common features or rules that govern the evolution and dynamics of an individual’s performance display. In all animal displays, audiences expect a form of authenticity or honesty in a performer. In animal behavior, this requires that a display conveys a more-or-less honest signal of Darwinian fitness. For example, in socially monogamous bird song, the male’s singing is an honest indicator of not only his physical quality but also the quality of resources that his defended territory may hold. Males on better territories can sing louder and more frequently than those on lesser territories [44]. In human singing, particularly in the genre of opera and other live musical performances, we seem to also respond directly to the ‘live’ presence of the performer [8] perhaps because we appreciate the physicality of the performance by responding well when performers give enhanced effort. And we reject forms of deception like lip-syncing in live performance [66], even though it is quite accepted, even necessary, in other forms of performance such as TV or music video. In the context of animal behavior, the cheating or falsification of specific single indicators of fitness often lead to the evolution of behavioral displays with many multi-variate or multi-modal components fitness. This is because a combination of feature components are harder to cheat as a whole than any single feature alone [15] [55]. This unwritten contract of authenticity between the performer and their audience also leads to increased variance and subsequent individuality among behavioral display across a population as well. The observation of increased variance in genetic, behavioral and/or morphological traits is one way evolutionary biologists have traditionally used to detect sexual selection acting on a species [67] [84] [79]. Similarly, in music, increased individuality and variance among professional performers can equate to a ‘signature sound’ that can be very essential for long-term mainstream succuss [25].

While compelling in its simplicity, a performance-based view of the function of music does not explain everything. For example, many forms of ‘ambient’ music have evolved where there is no apparent performer and the music seems to serve to create a space where the listener can enter and occupy. In this sense, perhaps ambient forms of music gives the listener the illusion of being in the center of a performance space as opposed to a seat in the audience. There is some evidence that this kind of music might also be less likely to stimulate the attention network of the brain and more likely to play upon self-awareness via the default mode network [41] [71] [63]. Music also has many cultural functions unrelated to individual performance that may have contributed to its evolution as well. These are mother-infant bonding (i.e. lullaby) [53], organization and bonding during prosocial enterprise (i.e. work songs) [74] and spatial-behavioral organization during military campaigns (i.e.. marching) [58]. These many culturally-defined aspects of music, perhaps layered over more ancient behavioral functions make it a challenging and complex subject of study.

Here we propose that some basic investigation of acoustical features of music that arouse attentive behavior in the listener could prove especially insightful regarding the common behavioral mechanism(s) that probably underlie the diverse cultural functions of music. The need for attentive behavior probably first arose during the transition from the Ediacaran to Cambrian (E-C) period about 541 million years ago when animal life first evolved swift directed movements as well as the more complex brains and sensory organs required to detect these movements. Associated with this E-C boundary are the rise of many new body forms, extensive burrowing behavior, and the evolution of protective shells covering soft body parts [16][56]. Attentiveness to swiftly moving bodies can be described mathematically by a simple calculus where an observer must accurately gauge the position of another body in time and space (Figure 1A). This requires accurate reading of the control of position of the other organism (f(x)), its energy or velocity relative to the observer (f’(x)), and its change of direction through acceleration and deceleration relative to the observer (f”(x)). An incomplete ability of the observer to perfectly account for changes in direction would create an element of surprise and unpredictability on the part of the observer. We can also similarly describe the quality of performance displays of an observed individual directly through their exhibition of motor control, kinetic or potential energy, and its cognitive reserve (i.e. ability to surprise). In the context of perhaps all behavioral displays of individual fitness across many animal species, including humans, we propose that this calculus of attention is always present to some degree, and that audiences of behavioral displays that represent individual quality will be attentively engaged by any components of the display that also convey honest signaling regarding motor control, energy expenditure, and surprise (i.e. complexity) (Figure 1B). We can directly relate this general idea to multi-modal tail display in peacocks (Pavo cristatus), where males perform a tail train rattling movements via feather stridulation at 25.6 Hz providing an audiovisual tail display where stationary eyespots are presented in front of a background of very rapid wave-like feather undulations to hold the attention of females and affect mate choice. [15]. In human music and dance, we can also readily observe elements of control, energy, and surprise (CES) and measure them in sound itself using a variety of audio processing methods (Figure 1C). We can mathematically define the CES of external sounds in a ternary space and map them dynamically over time (Figure 1D). We hypothesize that the more physically, energetically, and cognitively fit an individual musician or animal vocalist is, the better they can (A) control and stabilize their sound in the ternary space of CES and (B) move more intentionally within it (i.e. poor performance will create more randomness in the CES trajectory). We introduce an open-source, user-friendly software application named the POPSTAR PROJECT (https://gbabbitt.github.io/POPSTAR-signaling-in-music/) to dynamically map and analyze sound files (.wav or .mp3) in the space of CES. We introduce a stability indicator function for CES that measures song performance with respect to the calculus of attention. We then compare (A) musical and non-musical sounds, (B) various animal vocalizations, (C) musical genres with different emphasis on solo performance, and (D) differences in musical performance ability and environment (e.g. expert vs. novice, live vs. studio). We find compelling evidence that this ancient attention signaling is representative of an individual’s physical, energetic, and cognitive ability and is still retained in most modern musical performance today.

**Figure 1:**
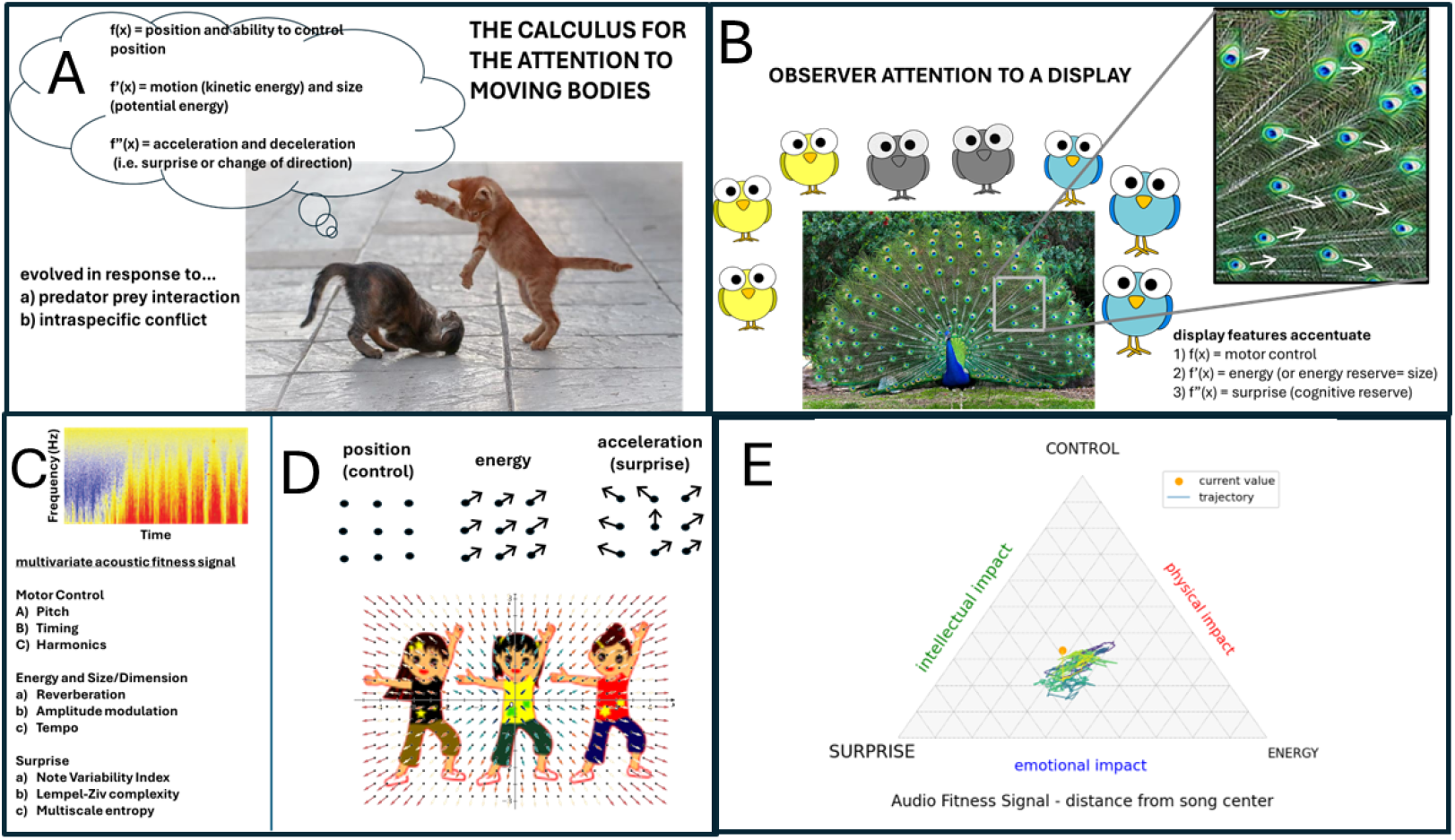
The theory of the calculus of attention examined through musical stimulation of human minds. (A) The calculus of attention focuses upon three components of biological fitness; motor control, energy level, and cognitive capacity demonstrated via surprise or novelty. (B) It is captured by many proximate stimuli present in animal displays. For example, a female bird might choose among prospective males based upon how loud or fast they sing (indicator of energy), how well they conform to a species-specific call (indicator of motor control), and in some species such as mockingbirds, nightingales, and lyrebirds, how well they can learn to improvise (indicator of cognitive reserve). (C) Human music and dance also capture attention using these same three elements of display. (D) These outward displays of attention signaling by an individual musician or group of musicians are plotted dynamically over time by the POPSTAR software and represented in a moving image ternary plot where the vertices are normalized multivariate measures of control, energy and surprise (likely processed via the dorsal attention network in the brain) and the sides combine to affect the internal focused physical, emotional, and intellectual sensations (likely processed via the default mode network of the brain). Example .mp4 videos can be found at our website https://gbabbitt.github.io/POPSTAR-signaling-in-music/

## 2 Methods

The POPSTAR software begins with the audio file in .mp4 or .wav format and segments it into user defined windows spaced on either (A) every beat using python’s librosa library beat tracker algorithm [20] (if user checks the box labeled ‘beats” on the GUI), or (B) every 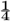 second if the user leaves ‘beats’ unchecked. We extract nine acoustic features of attention signaling from each audio segment, 3 each for control, energy, and surprise (i.e. CES). Features are extracted from a sliding window of user defined length that moves through the entire audio file. The default that is recommended by the software is a sliding window of 8s. We define performance quality directly in terms of acoustic measurement of physical and cognitive indicators of individual quality (i.e. fitness) such as motor control, energy, and complexity. Note that this is quite different than mathematical descriptions of Darwinian fitness as heritable phenotypic changes in populations (e.g. the Price equation [26]). While it is an obvious conjecture that direct indicators of physical and cognitive quality might translate to Darwinian fitness within populations, we do not measure or model this here.

### 2.1 Acoustic features of control (C)

#### 2.1.1 Control of pitch or frequency

Pitch control (PC) is defined as

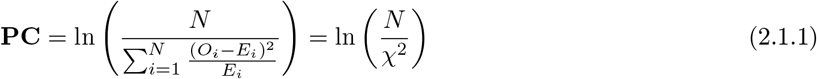

with

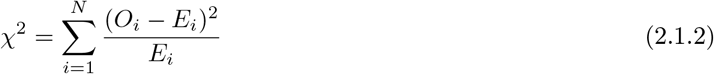

Where *N* is the number of notes discovered with a *f*_0_ or fundamental frequency and *O* = *f*_0_ in Hz determined by librosa pYIN algorithm [14][50] and E = expected frequency of the nearest value on the western music scale at A440 Hz or else is determined by nearest cluster mean frequency determined via expectation-maximization (EM) clustering. If the user chooses music’ or ‘single’ as sound-type on the GUI, then the Western music scale is used. If the user choses ‘other’ as sound-type on the GUI then the clustering approach is used.

### 2.2 Harmonics Control

Harmonics control (HC) is defined as the average harmonic energy (HE) across *N* notes, where each note contains *L* levels of harmonics detected above the fundamental frequency (*f*_0_) by the librosa harmonic spectrum analyzer [68]. The harmonic energy (HE) of a single note is the log-sum of energies *A*_*i*_ at each harmonic level *i*:

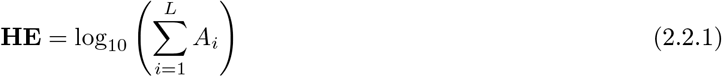

Harmonic control (HC) is the average of these energies across all *N* notes:

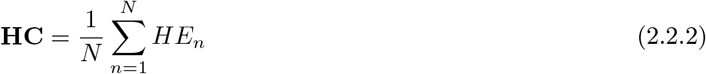

### 2.3 Timing Control

Timing control (TC) quantifies the stability of the beat interval. Let *O*_*i*_ be the observed beat interval and *E*_*i*_ be the expected (mean) beat interval returned by the librosa beat tracking software [20]. TC is defined as the log-inverse of the normalized mean squared error:

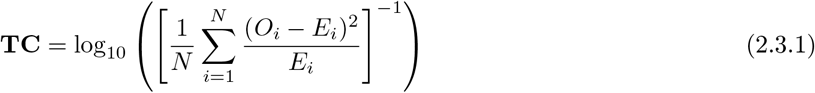

### 2.4 Acoustic Energy Features

#### 2.4.1 Tempo

Tempo (TP) is defined as the rate of beats over time:

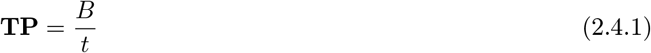

where *B* is the total number of beats and *t* is the time in minutes.

#### 2.4.2 Reverberation

Reverberation (RV) is determined by the autocorrelation coefficient at the first lag, representing the signal’s self-similarity:

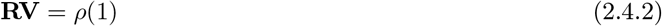

The Autocorrelation Function (ACF) at lag *k* is computed over the time domain signal *x* as:

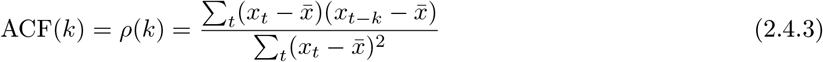

#### 2.4.3 Amplitude

The amplitude volume (AM) is determined as

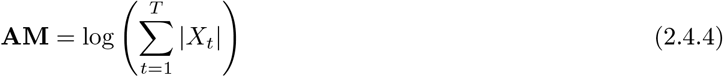

Where *X*_*t*_ is the audio signal in time domain and *T* is the time length of the audio segment

### 2.5 Acoustic features of surprise (S)

#### 2.5.1 Multi-scale entropy

The multi-scale entropy (MSE) is defined as the complexity index returned from the refined multi-scale entropy (rMSE) function for univariate data from python’s EntropyHub library [24] using an Entropy object with embedding dimension =4 and radius threshold = 1.25 to return a Complexity Index over 5 temporal scales using 3rd order Butterworth Filter with normalized corner frequency of 0.6 at each temporal scale.

#### 2.5.2 Lempel-Ziv complexity

The Lempel-Ziv complexity (LZC) is defined as

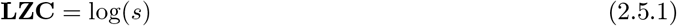

Where s = number of distinct substrings in the binary representation of the audio file

#### 2.5.3 Note variability index

Note variability is defined according to the method of [77] for analyzing bird song complexity.

The spectral cross-correlation is converted to a Note Variability Index (NVI), ranging between 0 and 1, where:

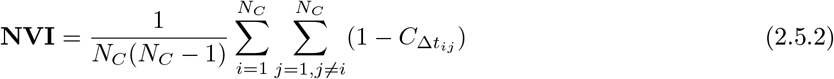

This forms an *N*_*C*_ × *N*_*C*_ similarity matrix, where *N*_*C*_ is the number of notes and 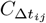 is the association between notes *i* and *j*. Thus, a lack of cross-correlation across all pairs of notes or data columns in a spectrogram represents high variability in the sound file while a tone repeated constantly in the time domain has high cross-correlation but no complexity. It is important to understand that variability in the signal increases to the point of unpredictability, thus the variability eventually increases from a complex pattern to memoryless white noise (where the next value in a sequence cannot be predicted at all).

### 2.6 Min max (self) and Z score normalized average values of 9 features of CES

If the user chooses ‘single’ file under sound-type on the GUI, the CES feature matrix constructed listed above is min-max normalized by column where

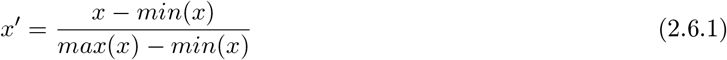

If the user chooses ‘music’ under sound-type, the CES feature matrix is z-score normalizes using the average feature parameter values (*µ, σ*) for human music where

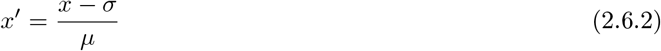

If the user chooses ‘other’ under sound-type, the CES feature matrix is z-score normalizes using the average feature values (*µ, σ*) for human speech.

### 2.7 Dynamic ternary plot representation of CES over time

Control, energy and surprise features for each audio segment in time (t) are averages of the normalized features from that audio segment

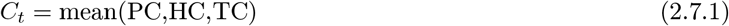

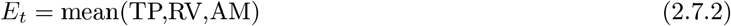

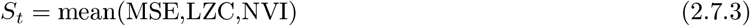

and thus

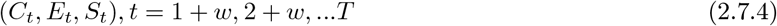

where t is user defined as 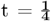 second OR beats tracked in the time of the music and w = analysis widow length determined by user (default is 8 seconds) and values of (CES)t are normalized to sum to 1

The single points in CES ternary space are thus

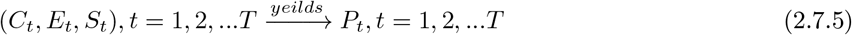

The computational workflow and plate notation for the operational model used here in the POPSTAR software is presented in Figure 2.

**Figure 2:**
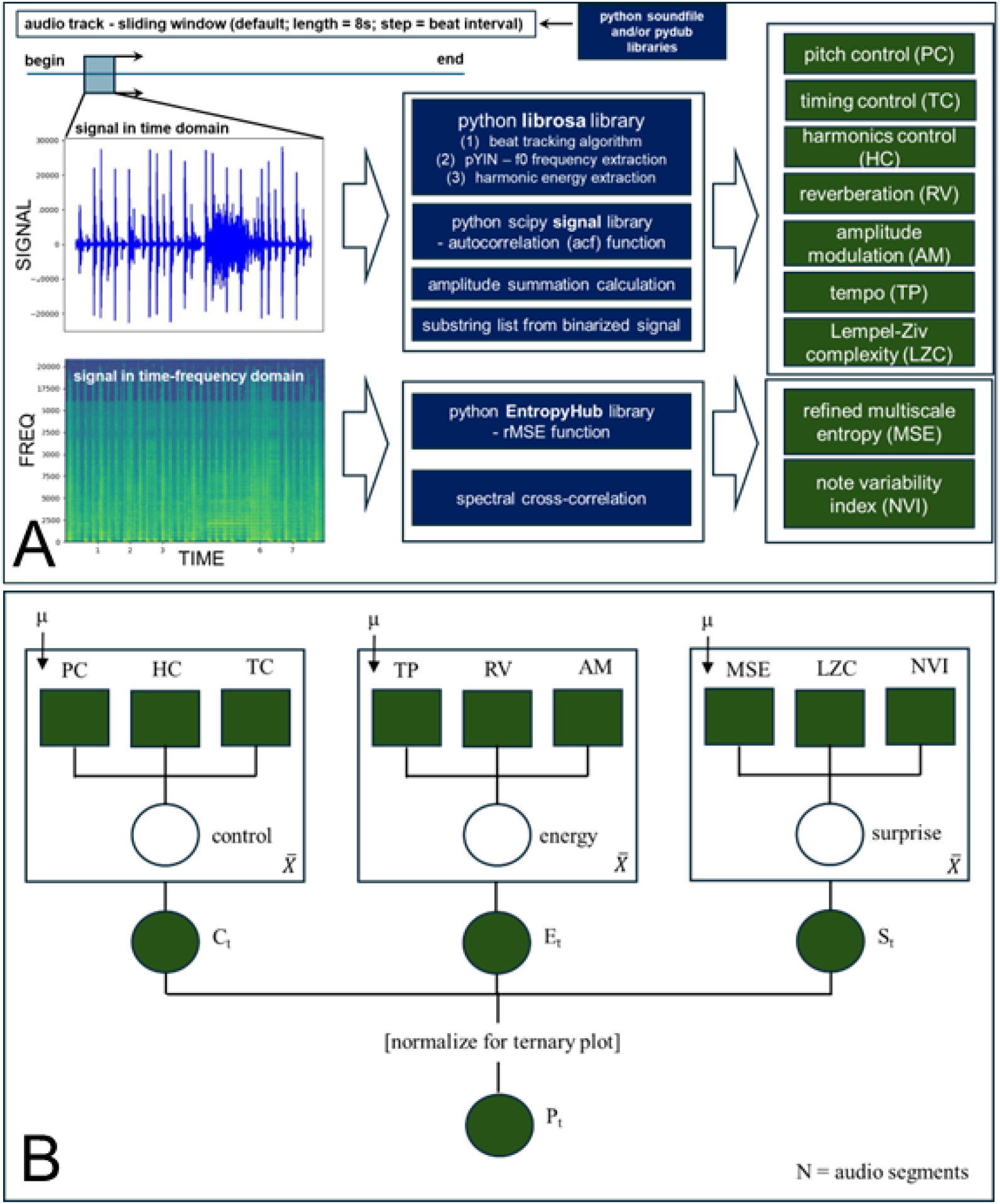
The computational workflow diagram for the POPSTAR software and plate notation of the multi-modal model analyzed within. (A) The POPSTAR software uses a sliding window to discretize the input audio file into segments of equal length that are preprocessed via the python soundfile and pydub libraries to create time domain and time-frequency domain representations of each audio segment. These are post-processed via python librosa and our own custom functions to extract 9 measured features; 3 each representing the latent variables of control, energy, and surprise (i.e. complexity). (B) A plate notation of the model that converts the 9 features into a single point in ternary space is also shown. Here squares represent the inputs to the model and circles represent outputs. Latent variables are left unshaded while observed variables are shaded.

A dynamic ternary plot is generated as well as a dynamic Chernoff face [13] and saved in movie file (.mp4) format (Figure S1). In the Chernoff face representation, Eye height = NVI, Eye width = LZC, Pupil area = MSE, Eyebrow slant level = TP, Mouth height = AM, Mouth width = RV, Ear height = PC, Ear width = TC, and Nose slant = HC. Examples of .mp4 (movies) containing dynamic Chernoff faces and dynamic ternary plots with sound overlay for a variety of bird song, human/animal vocalization, and human music can be found in examples dynamicPlots.zip. Thus, the POPSTAR software produces a dynamic Chernoff face using all 9 features tracked over time where the three features of control are represented by the ear dimensions (and nose angle), the three features of energy are represented by the mouth dimensions (and eyebrow slant), and the 3 features of surprise are represented by the eye dimensions. It has been observed that human minds struggle to conceptualize statistical relationships that extend much beyond 3-dimensional space and have trouble tracking more than a handful of objects in real time [12]. Herman Chernoff invented facial representation of complex data because faces are one of the only multivariate structures that evolution has equipped us to rapidly and easily interpret [13]. The normalized averages of the control, energy and surprise are also plotted dynamically as a song trajectory within a ternary plot. See the POPSTAR software landing website and Figure S1 for more details.

### 2.8 Stability indicator (i.e. performance fitness indicator) for the tracking of CES over time

We develop a simple permutation-based indicator of the stability of the performer’s trajectory within the space of CES (Figure 3). It can be observed that both animal vocalizations broadly characterized across anuran, avian, and primate taxa have histograms where most steps in the ternary space of CES are quite small. This distribution tails off with a power lognormal decay often with some multimodal behavior observed in the tail (Figure 3A). Human music is quite similar with even more stability in CES step length (Figure 3B). We hypothesize that high levels of performance quality and resulting musicality of sound (i.e. fitness of performance) should result in more stable and intentional movement within the ternary space of CES. To measure the stability of any given audio signal we create permutations of the observed CES trajectory by randomly shuffling time steps and comparing the resulting observed and randomized distributions of step length in the ternary space of CES (Figure 3C-D). Note this does not alter the values of points in the ternary space, only the order in which they occur, thus

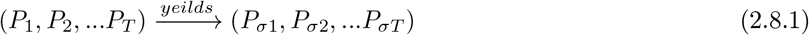

**Figure 3:**
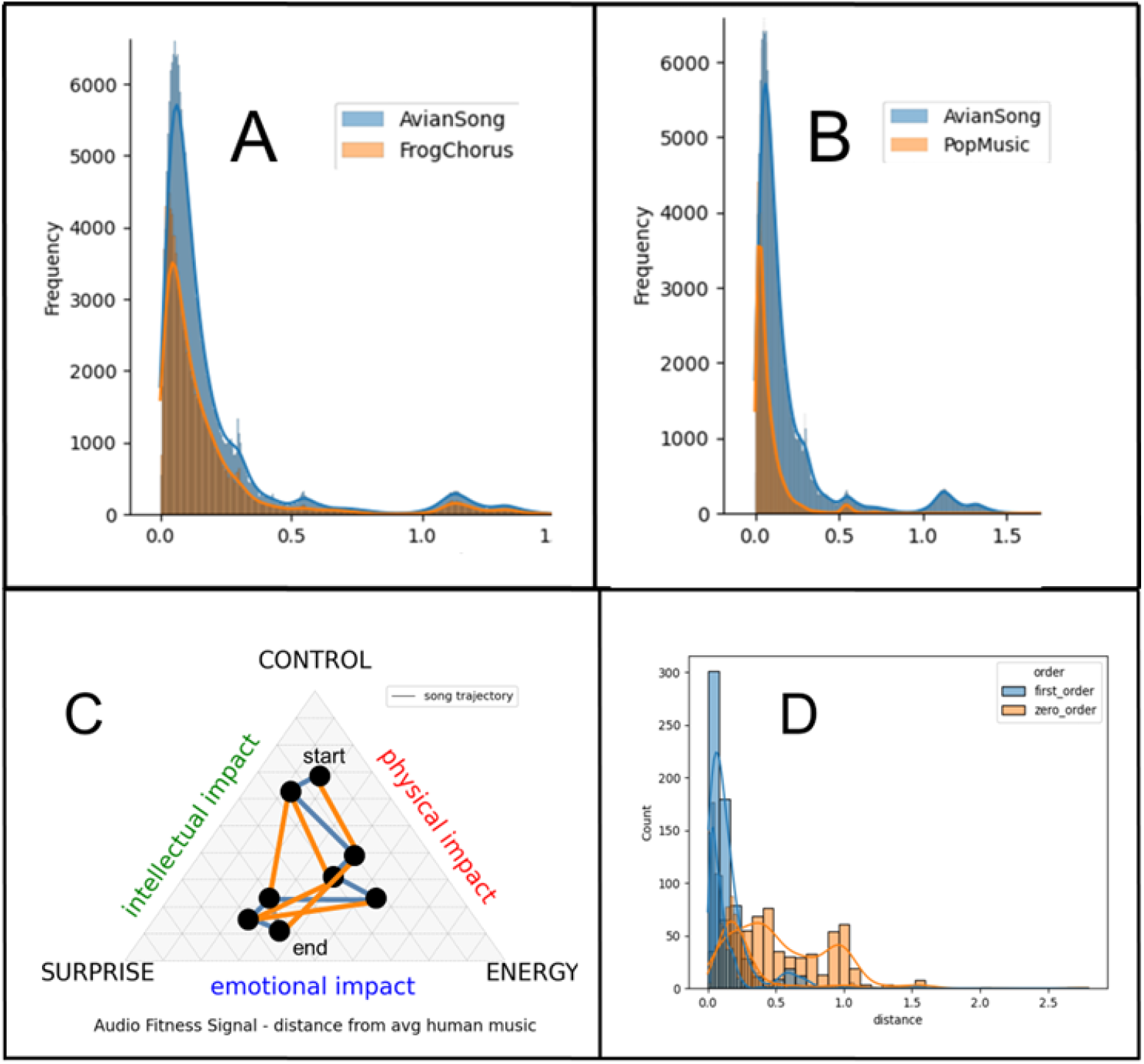
**The shape of the distribution of step distances in the ternary space of control, energy, and surprise (CES) in sound** demonstrates a strikingly common pattern across (A) anuran chorus, avian song that is even more stable in professional (B) human musicians. (C-D) The stability of song trajectory in the space of CES is determined by comparing the step distribution of the song trajectory in the space of CES (blue) to random walks of the song trajectory created by the shuffling of the order of steps (yellow). An expertly created musical sound typically shows a multi-modal lognormal, gamma or power lognormal distribution of steps where step distances are usually very short and controlled until the musician intends to move somewhere else in the space of CES. When the time dependence of the signal is destroyed by shuffling (C), the distribution of steps (D) becomes more uniform with a larger mean step distance. In the case of brown noise (not shown), no difference in observed and shuffled step distributions is evident. The percentage of the stability in CES can be determined through repeated permutations of random draws from the real- and shuffled-time step distributions and tallying the number of times the time step in the observed trajectory is less than the that of the shuffled trajectory. The significance of this difference can be determined via the Kruskal-Wallis test.

Each observed time step in the ternary space

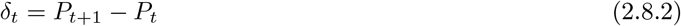

is compared to each corresponding shuffled (i.e. random) time step

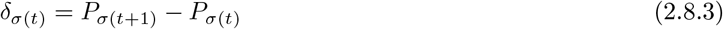

and we count instances in which the observed step is less than or greater than its corresponding shuffled step

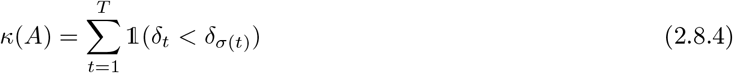

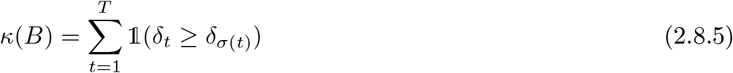

Thus. an indicator of stability ranging from 0 to 100 percent in the CES trajectory becomes

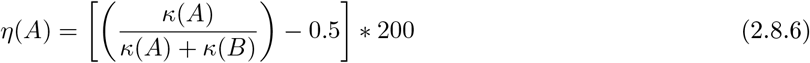

Nonparametric hypothesis testing on average *C*_*t*_, *E*_*t*_, *S*_*t*_ and stability *η*(*A*) can be achieved using the Kruskal-Wallis test where the test statistic *H* is given by

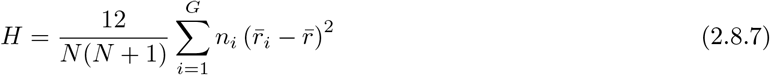

Where *N* is total observations across all groups, *G* is number of groups, *n*_*i*_ is number of observations in group *i, r*_*ij*_ is rank of observation *j* from group *i*, 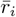 is average rank of all observations in group *i*, and 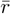 is average (grand mean) of all the *r*_*ij*_’s. Multi-model inference can also be performed on the observed step distributions using Bayesian information criterion

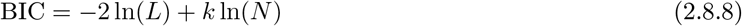

where *L* is likelihood of model given the data, *k* is the number of parameters in the model and *N* is the number of data points

### 2.9 Comparative functional data analysis of song trajectory in the dimensions of control, energy, and surprise

For comparative analysis of different groups of songs (i.e. human artists, musical genres, characteristic species songs, etc), we fit b-spline functions to *C*_*t*_, *E*_*t*_, *S*_*t*_ trajectories of each song using an optimal tuned bandwidth determined via the highest group classification accuracy achieved when employing the K nearest-neighbors classifier function from python scikit-fda module. The time-series functions were aligned (registered) using Fisher-Rao elastic registration [82] from python scikit-fda applied to all songs in the groups being compared. Group machine learning classifications were then conducted using four methods in scikit-fda (KNN, nearest centroid classifier, maximum depth classifier, and a functional quadratic discriminant analysis classifier). Overall classification performance was determined using these four methods as a multi-agent stacked model (Table 1). The individuality or dissimilarity between any two songs was defined as the *L*^2^ or Euclidian distance between the elastic registered b-spline functions for each song in each of the three dimensions *C*_*t*_, *E*_*t*_, *S*_*t*_. An adjacency matrix was constructed for all pairwise comparisons between songs and represented as a Kamada-Kawai force directed graph network [40] with edges constructed between allindividual *L*^2^ distance that fall below the median of all *L*^2^ distances. Thus the 50 percent closest nodes in the network graph have an attractive force and the highest 50 percent have a repulsive force and the network is therefore well balanced in its modularity. Graphs and modularity calculations of the network was computed using python networkx. See Figure 4 for a schematic example of this.

**Table 1:**
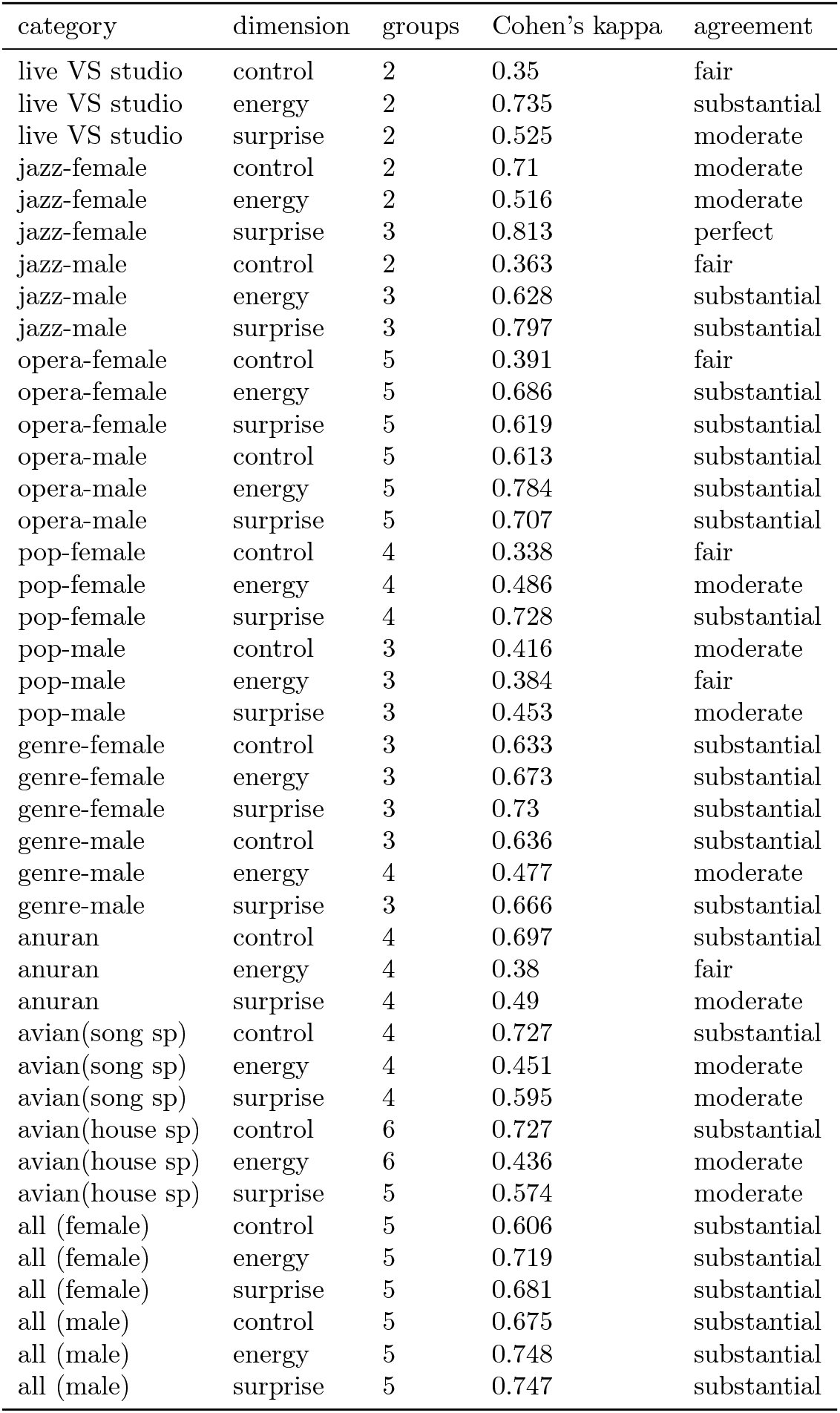
Overall agreement for the FDA machine learning classification tasks with the various categories comparisons in our research. Classification was performed with a multi-agent model including a maximum depth, nearest centroid, K nearest neighbor and functional quadratic discriminant analysis. Model accuracy was converted to Cohen’s kappa based upon the number of grouping factors it was deployed upon. The live vs studio comparison involved only the popular musician Bjork. The musical genres compared were popular music, jazz and opera. The anuran (chorus frog) comparisons included 4 species from the genera Hyla and Pseudacris. The avian comparisons included Nightingale, Superb Lyrebird, Northern Mockingbird, Brown Thrasher, Catbird, and either Song Sparrow or House Sparrow. The comparison of all animal groups included the anuran, avian, opera, jazz, and popular musicians combined within each group.

**Figure 4:**
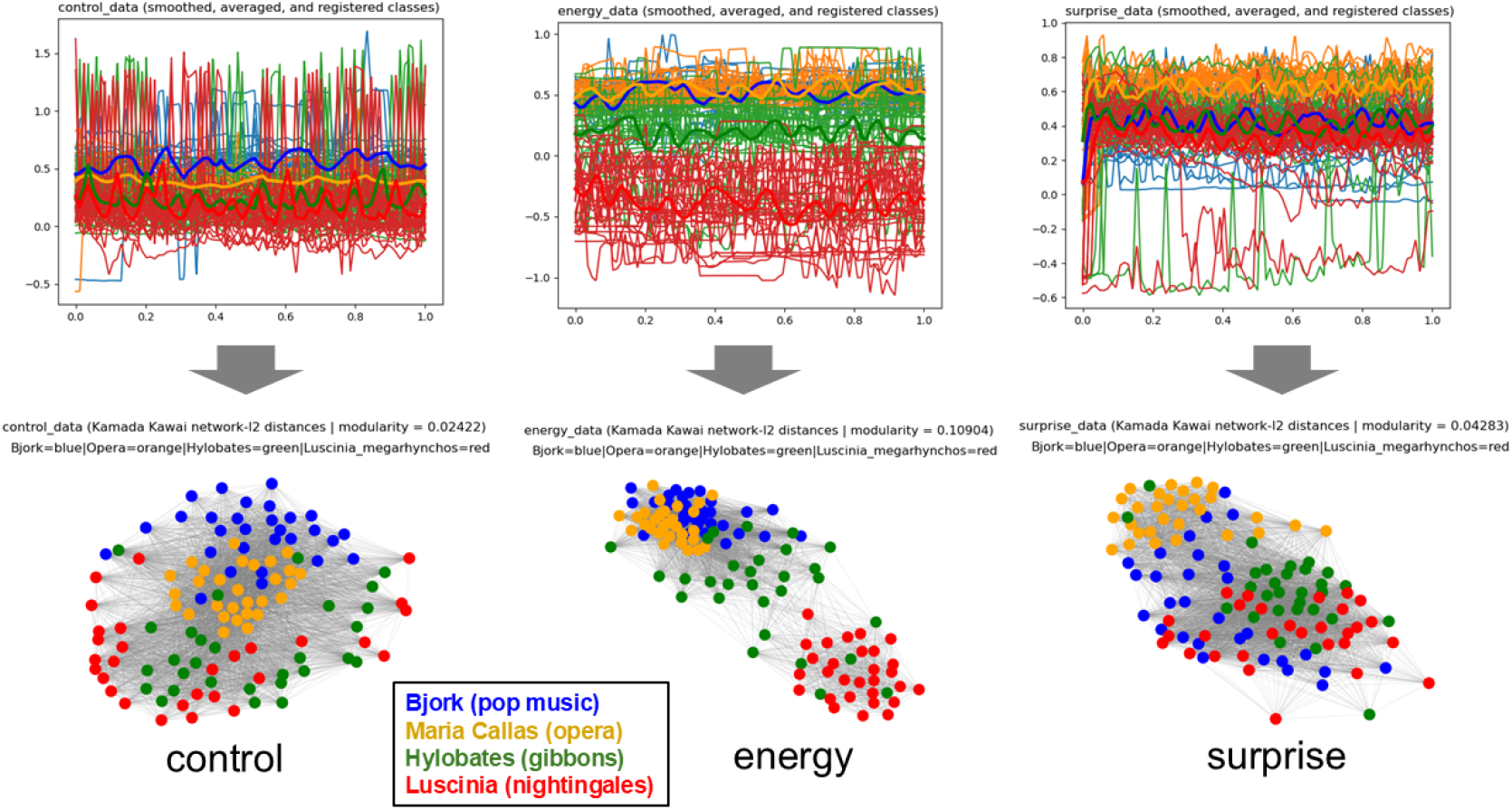
Functional Data Analysis (FDA) and adjacency networks for comparatively analyzing the similarity/individuality of single song time series trajectories in the space of control, energy, and surprise (CES). First b-spline functions are generated and aligned using optimal bandwidths determined via machine learning classification of the grouping classes after 70/30 training-test split (method = nearest centroid classifier from python scikit-fda). L2 or Euclidian distances are calculated between all pairs of spline functions in the FDA to create an adjacency matrix. The network is mapped using the Kamada-Kawai method (python – network module) with edges placed between all L2 distances below the median L2 value.

### 2.10 Random forest classification

The 9 normalized acoustic features are used to train subsequent random forest models that were used to assess comparative classification of sound types derived from our protein sonification when compared within the context of human speech, music, and other natural and computer-generated sounds. All sound files used are listed in folder list.txt and included in Supplemental Data. Note that some music files are omitted due to copyright, but the titles of the music tracks analyzed can be seen in the first column of featureData allFolders.dat. Our random forests consisted of 500 decision trees using 80/20 percent training/testing data split. It is implemented with python scikit-learn RandomForestClassifier function which implements the ID3 with CART algorithm to create each decision tree.

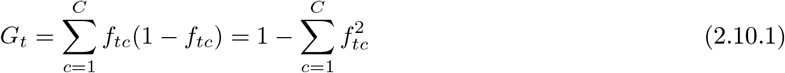

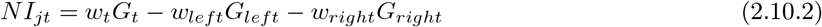

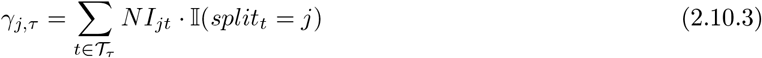

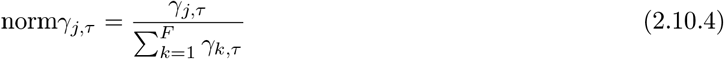

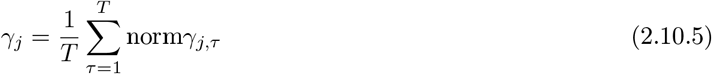

### 2.11 Data retrieval and statistical comparisons

In our research, we use the POPSTAR project code, popstar.py available at https://gbabbitt.github.io/POPSTAR-signaling-in-music/ to generate dynamic ternary maps and make statistical comparisons of CES and attention signaling (i.e. CES stability) across musical and non-musical sounds, musical genres, and several artist’s discography, and selected examples of human, primate, bird, and anuran(frog) vocalizations. All animal vocalizations were extracted from the iNaturalist dataset() audio repository iNat sounds available at https://github.com/visipedia/inat_sounds/ All iNaturalist audio files were manually curated for consistency of song type and cleanness of audio, removing any with background sounds of humans, other animals and/or excessive wind noise. Movie output for specific examples of the dynamic ternary plots and dynamic Chernoff faces can be observed at the website above. We make specific comparisons of attention signaling in expert vs. novice vocalists in both humans and Yellow Canary (owners often post the first songs of their pet canaries on YouTube after training them with videos of ‘exceptional’ adult birds singing). We compare attention signaling in piano concertos, where the ability of soloists is key, to ambient piano works of the Impressionist era, where the music emphasizes the environment of the listener rather than the skill of the soloist. We also compare live vs. studio performances of the popular artist Bjork Guomundsdottir, to see if direct feedback of the audience affects the resulting attention signaling of the artist.

In cases where audio was extracted from YouTube, we avoided videos with excessive background noise and performed batch audio extraction and batch loudness normalization using our script **(yt-dlp batch.py)** available in the POPSTAR Project Github repository. All ternary plot movie output (with sound) and statistical results are available at https://doi.org/10.5281/zenodo.17218769.

## 3 Results

### 3.1 Validation of musicality of sound via stability of control, energy, and surprise

In comparisons of stability of audio trajectory in the ternary space of CES (i.e. signaling of individual performance quality) across a large variety of musical and non-musical sounds (Figure 5), we find that music has the highest stability in CES (>60-70 percent) while samples of brown noise have near 0 percent stability (Figure 5A). Therefore, our indicator function for stability in CES is a valid measure of the human perception of musicality in sound. We find that famous human musicians in the genres of jazz, opera, and popular music have significantly different and higher CES trajectory stability than other animal vocalizations (H = 416.945, p <0.001) with some birds with very complex song such as the Lyrebird of Australia (Menura novaehollandiae) and the European Nightingale (Luscinia megarhynchos) having CES stability that is comparable or even surpassing some human singers (Figure 5B). We also find that the career discography of a single pop musician (Bjork Guomundsdottir) also exhibits significant variation in CES stability (H = 85.492, p< 0.001) (Figure S2) indicating that our metric of musicality is not only valid regarding our perceptions of song, but has potentially interesting variation within individual performances as well.

**Figure 5:**
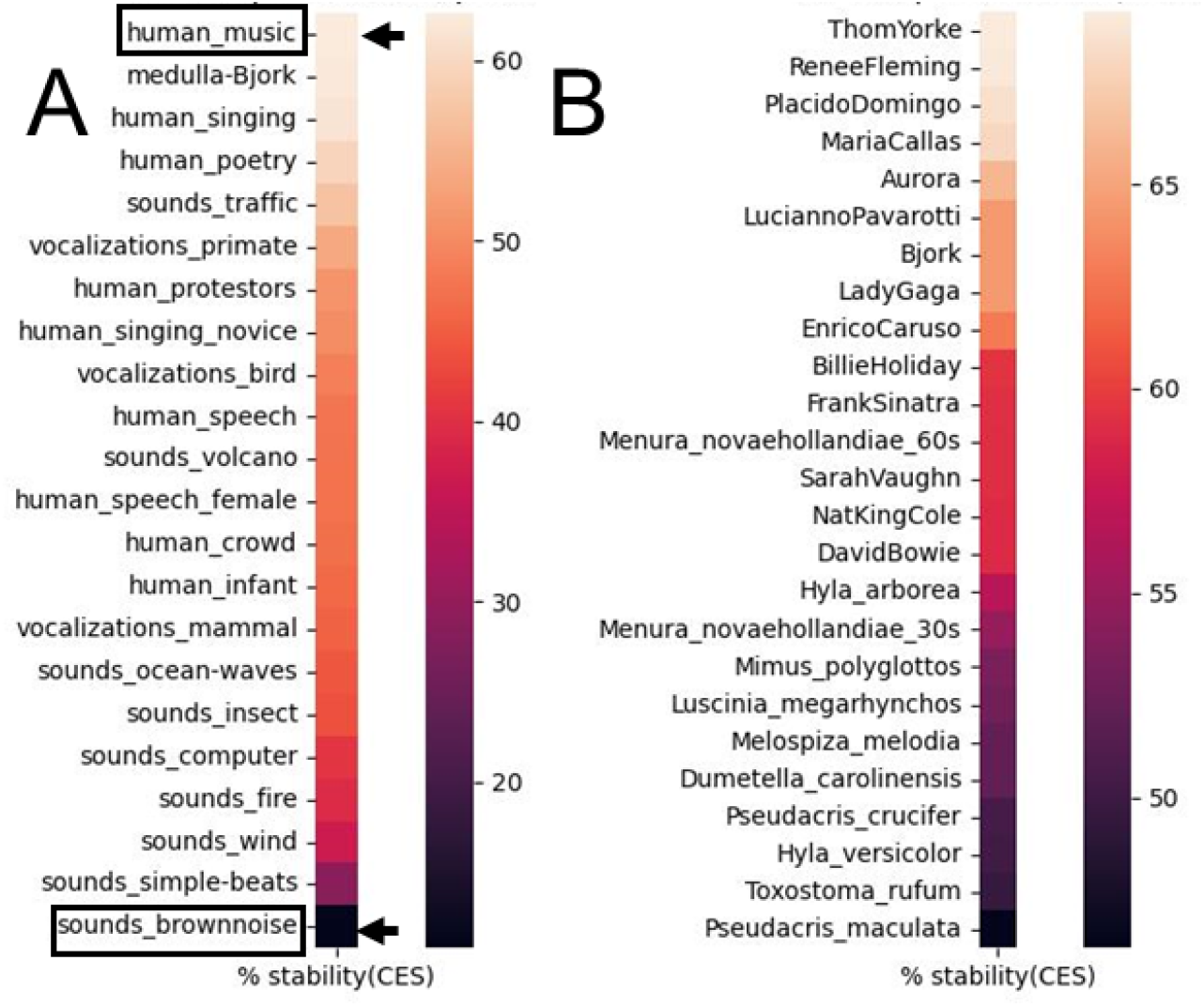
**Relative stability of different sound trajectories in the space of control, energy, and surprise (CES)** for (A) a selection of natural sounds ranging from noise, non-biological sounds in nature, and various animal (and human) vocalizations including music. (B) We also compare a selection of famous human musicians of various genres (Pop, jazz and opera) to chorus frogs (genus Hyla and Pseudacris) and several birds with highly notable songs (i.e. Song Sparrow (Melospiza melodia), Lyrebird (Menura novaehollandiae), Nightingale (Luscinia megarhynchos), and several species of Mimidae (i.e. mimics) including Northern Mockingbird (Mimus polyglottos), Brown Thrasher (Toxostoma rufum) and Catbird (Dumetella carolinensis). Note that (A) noise has 0 percent stability in the space of CES while music has the most (typically > 50 percent) which is roughly 10 percent higher than spoken words.

### 3.2 Comparison of control, energy, and surprise in human vs. animal song

A broad taxonomic comparison of individuality in song across humans, birds and chorus frogs across each dimension of CES (Figure 6) reveals that while professional human singers have less variation in control and energy than these two animal groups, the human genres are quite discernable from each other with opera being closer to animal vocalization in terms of energy and control than other musical genres. We also see that opera is closer to anuran than avian song (Figure 6A-B, 6D-E) perhaps due to more similar vocal architecture between mammalian and anuran groups, both having a larynx as opposed to the avian syrinx. In anurans, the larynx is used within a closed cycle air space moving air repeatedly over the vocal chords between a storage sac and the lungs. In mammals, air is inhaled and exhaled over the vocal chords. In birds, this air cycling is similar to mammals, but the avian syrinx sits deeper in the lungs with vocal chords on each side that can often be controlled independently allowing for more complexity of sound. In the dimension of surprise (Figure 6C, 6F), we still see differentiation between human singers of different genres and our two main animal groups, but the variability between human singers is more pronounced indicating more human variation in song complexity than with control and energy. This is perhaps not surprising given the language components of human song as well as the falsifiability of energy and control signaling in human music via the use of instrumentation and other technology and professional training. We do not observe much difference in these trends based upon whether the human singers are female (Figure 6A-C) or male (Figure 6D-F), however we do note that female human singers have somewhat more individual variation in song control and energy, while male human singers have more variation in song complexity or surprise. These trends are still apparent when we include a limited number of other ape songs (i.e. Hylobates – gibbons) in the analysis (Figure S3) and we note that primate song is also generally closer to humans as one might expect. Within chorus frog and avian species (Figure S4 and S5) we observe less individuality than across major taxonomic groups, but modularity in the song networks is still apparent especially with regards to control and surprise in anurans and energy and surprise in birds. Within human musical genres (Figure S6) opera and jazz tend to be more distinct emphasizing surprise (jazz) and energy+surprise (opera) while popular music emphasizes energy+control, with popular music having wider variation in CES distances between individual songs. This is perhaps not surprising given that popular music borrows from many other genres and thus may appear less distinct in our network representations. Within each musical genre (Figures S7-S9), while significant differences in the levels of CES are evident, we find little difference in the modularity of the network distances (i.e. individuality of each song).

**Figure 6:**
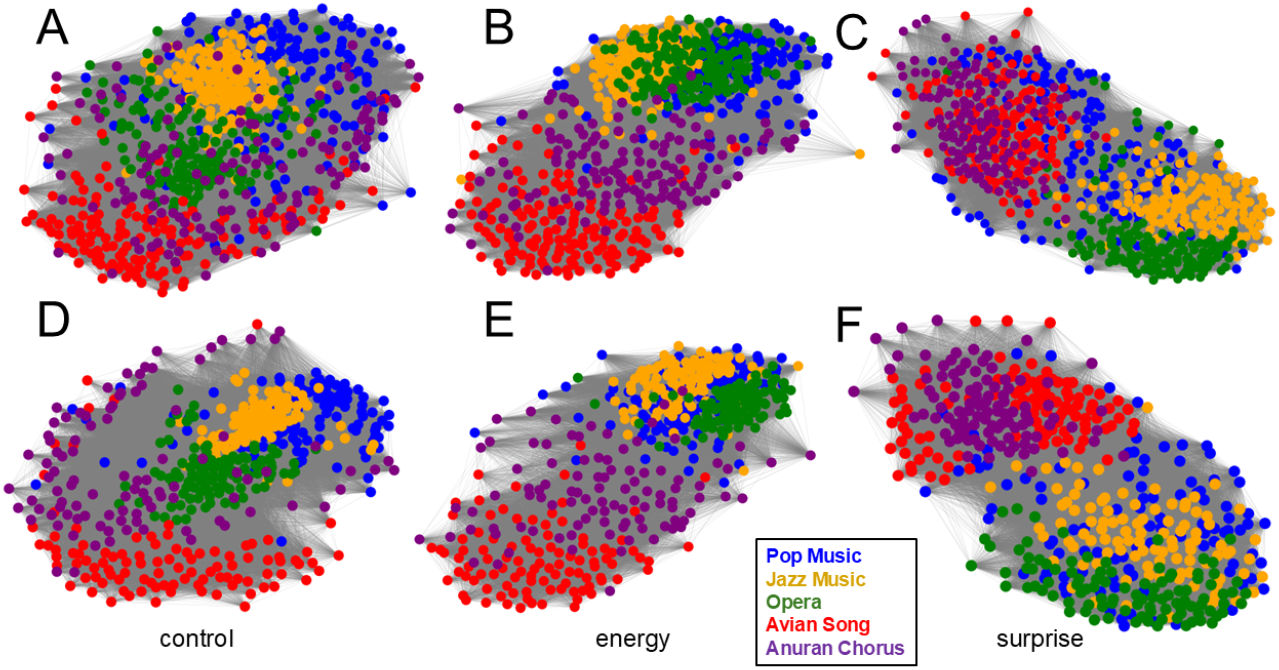
Adjacency networks of individual songs of frogs, birds, and humans mapped within the three dimensions of control, energy and surprise (CES). In A-C the human singers are female and in D-F they are male. In the animals, all singers are assumed to be male. Functional data analysis is applied to generate a b-spline interpolation of the time series of an individual song in the specified dimension (either C, E or S). In each network, each node represents an individual song and each edge represents an L2 distance between the functions for two songs. The network representations utilize Kamada-Kawai topology with edges formed using all L2 distances below the median distance observed (i.e. closest 50 percent of the nodes attract and the farthest 50 percent of the nodes repulse).

### 3.3 The role of audience feedback on control, energy, and surprise

We analyze the career of the pop musician Bjork who is very interesting from the context of our research because of the large range of ‘controlled’ experimentation that she has recorded. Across a nearly 50-year career and 6 World tours, she is well known to be an excellent live vocalist performer and as well a highly experimental composer touching upon many unique genres (pop, punk, trip-hop, jazz, folk, electronica, ambient, and avant-garde). Her vocal exploration of the boundary between her own human speech and her more conventional music is also important for our study, particularly her fifth studio album Medulla, which attempts to produce fully modern music using only the human voice alone (i.e. no instrumentation) (Figure S10). After each of her world tours she released a live album corresponding with the same tracks as her studio work prior to touring, thus providing a nicely controlled pairwise experiment at the level of each album and song where we can examine the effect of the feedback of live audiences on the different components of attention signaling in her performances (Figure 7). While we find no significant differences between live and studio performance regarding the stability of CES trajectories, we do see different levels of CES evident across 6 studio LPs recorded over 18 years (Figure 7A) that are absent in the corresponding live performance LPs (Figure 7B). This indicates more consistent combination of control, energy and surprise when facing a live audience. Combined comparison of live vs studio LP’s (Figure 7C) finds significantly increased control (H=11.918, p=0.001) and decreased surprise (H= 8.194, p=0.004) again indicative that audience feedback greatly effects attention signaling in a musical performer. We also note that the differences between live vs. studio performance is not reliant on any single feature of CES (Figure 7D) and is quite apparent in our network analyses of individuality of songs (Figure 7E). Lastly, we compared Bjork’s attention signaling across normal speech (i.e. TV interviews), her Medulla LP, and her Greatest Hits LP (Figure S10) and find significant differences in all levels of CES (control: H=39.034, p<0.001 | energy: H=15.014, p=0.002 | surprise: H=26.961, p<0.001) and stability of CES (H=16.033,p=0.003) with clear evidence from the Medulla LP indicating that the attention signaling present in the audio is not dependent upon electronic instrumentation, but can be the product of human vocalization alone.

**Figure 7:**
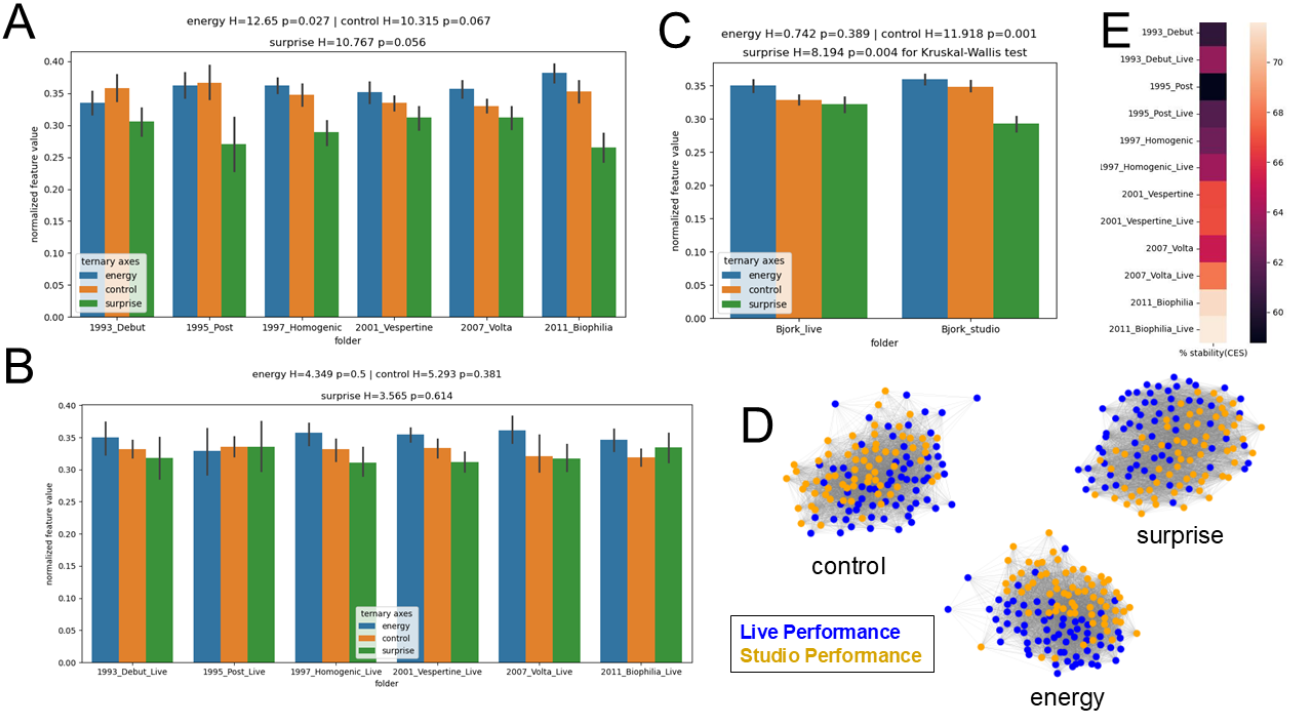
Comparison of stability in musical control, energy and surprise (CES) in live vs. studio performances while controlling for individual song track. Here we examine the Icelandic pop star Bjork’s six studio LPs compared to their corresponding live tour LPs. We find that (A) the studio LP’s exhibit significant or borderline significant differences in CES while (B) the corresponding live LP’s do not. This suggests that live performances feedback may constrain Bjork’s performance to a more consistent level of relative CES. (C) Overall, the live performances show significantly less control and more surprise than the corresponding studio song tracks, showing a trend similar to that distinguishing novice from expert performance. This suggests that fitness signaling via CES is less manipulatable on stage than in studio. (D) Random forest classification indicates with moderate accuracy that differences between Bjork’s live and studio performance involves all nine features of CES to large extent. (E) While each studio project and its accompanying tour vary significantly in CES stability, the live performances seem to be slightly more stable in CES, again suggesting some creative constraints or added fitness requirements when performing live.

### 3.4 Comparison of control, energy, and surprise in expert vs. novice singers

Comparisons of expert and novice human singers (Figure 8A-C) reveal that expert musicians exhibit borderline significantly more CES stability (H=4.339, p=0.037), more control (H= 3.429, p=0.064) and less surprise (H=5.357, p=0.021) than novice musicians (Figure 8A-B). A machine learning random forest classifier identifies expert vs. novice human singers with 99 percent accuracy using mostly the features of Lempel-Ziv (substring) complexity and note variability and amplitude modulation to a lesser extent (Figure 8C). In a similar analysis comparing adult Yellow Canary to immature birds singing their first songs (Figure 8D-F, Figure S11) we find no differences in overall levels of control, energy or surprise, but we do find that adults have significantly more musicality indicated by the stability in CES (H=3.971, p=0.046)(Figure 8E). Machine learning classification can distinguish adult from immature song with 93 percent accuracy using a wide variety of features of CES (Figure 8F).

**Figure 8:**
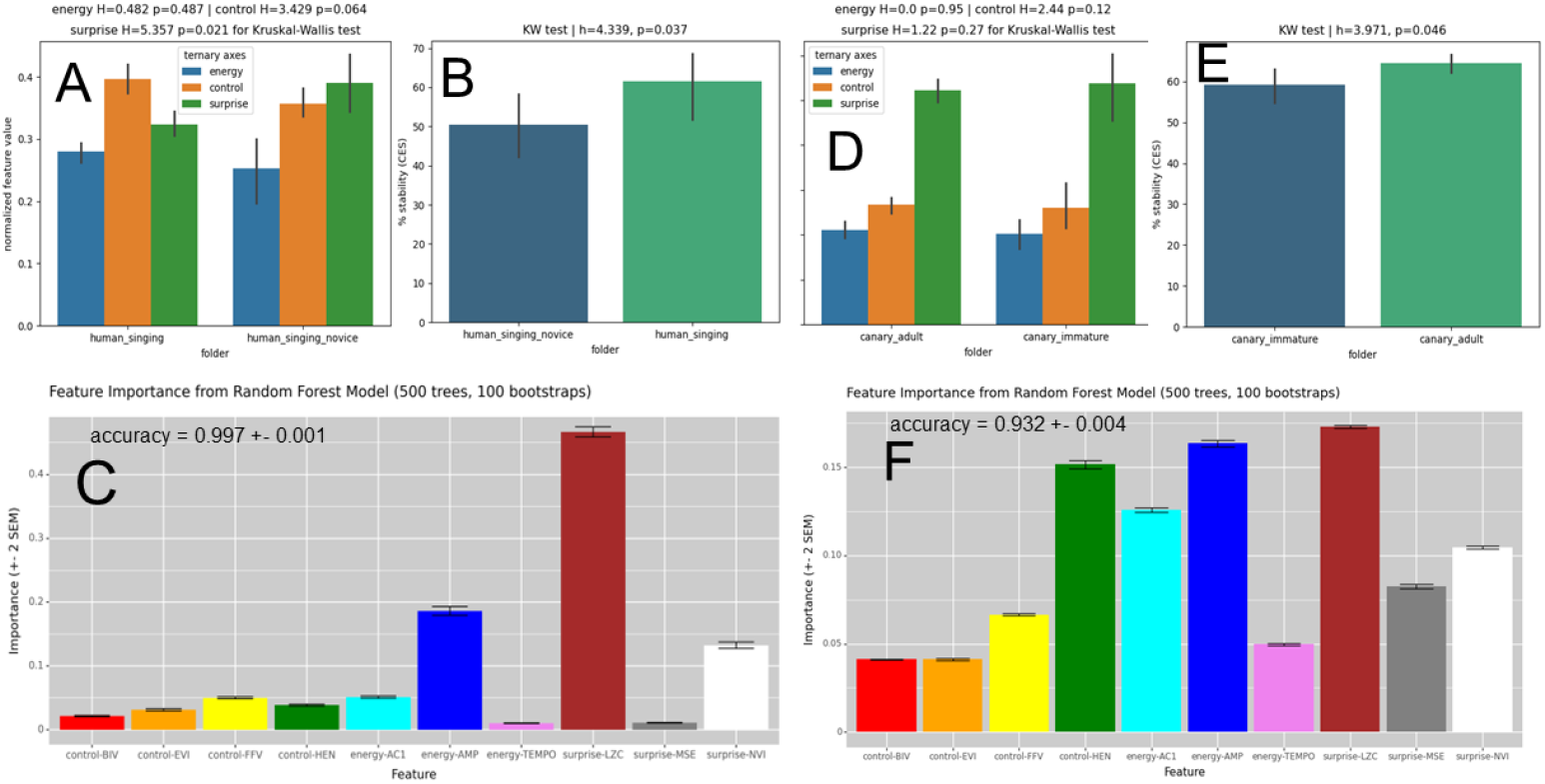
Comparison of stability in musical control, energy and surprise (CES) in expert vs. novice singers (A-C) and adult vs. immature birds (D-F). (A) Expert singers exhibit significantly more control and less surprise or complexity than novice singers. (B) Expert singers also have significantly more stability in song trajectory in the space of CES than novice singers. (C) Random forest classification indicates with high accuracy that differences between experts and novices are largely driven by Lempel-Ziv (substring) complexity, amplitude modulation, and note variability in this study.

### 3.5 Comparison of control, energy, and surprise in piano concerto vs. ambient works for piano

As not all music places equal emphasis on solo performance (i.e. individual performance quality), we compared a small sample of the most difficult piano concertos for soloists to perform (Rachmaninov Nos. 2-3, Beethoven No. 5, Mozart Nos. 20-21, Brahms No. 1, Tchaikovsky No. 1, Grieg Am concerto) with a small sample of ambient piano pieces by the composers Satie (Gymnopedie 1-3, Gnossienne 1-6), and Debussy (Arabesque No. 1-2, Clair de lune, Reverie) (Figure 9). Here we find that while the levels of control energy and surprise are similar (Figure 9A), the stability in CES is significantly higher in the concertos designed to demonstrate soloist virtuosity than in the ambient pieces which emphasize music as a space which the listener occupies (H= 5.357, p=0.021) (Figure 9B). Random forest classification can discriminate concertos from ambient works on piano with 95 percent accuracy indicating that multiscale entropy and reverberation are the most important features for learning the classification here (Figure 9C).

**Figure 9:**
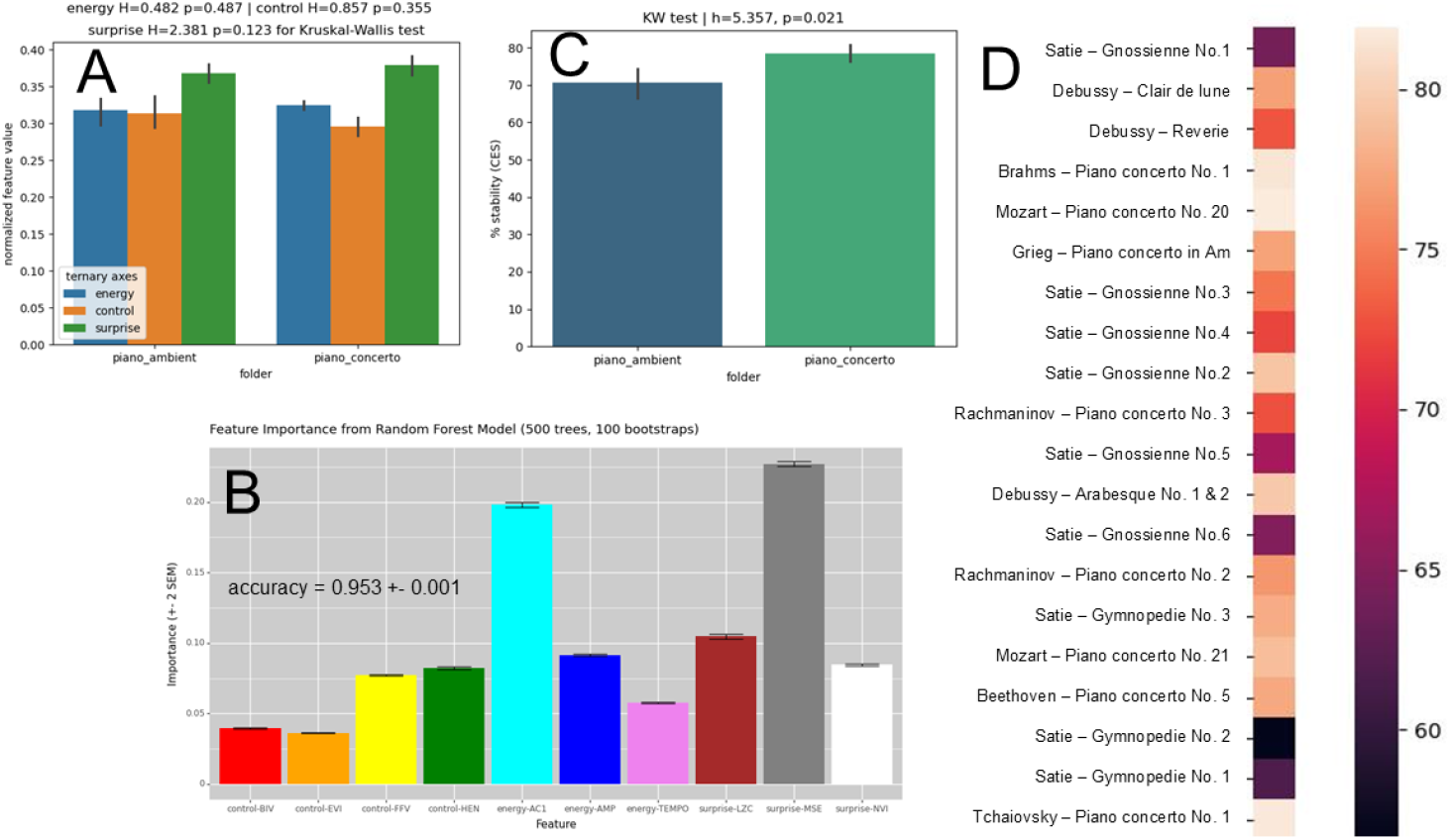
**Comparison of stability in musical control, energy and surprise (CES) in piano concertos vs. ambient piano pieces** (i.e. furniture music of Erik Satie and Claude Debussy). (A) While there are no significant differences in the levels of CES between the two genres, (B) the piano concertos have significantly more stability in song trajectory in the space of CES than ambient music. (C) Random forest classification indicates with high accuracy that differences between the two genres are largely driven by multiscale entropy and reverberation.

## 4 Discussion

We have proposed a novel multivariate measure of vocal performance (i.e. individual quality) in sound defined directly at the phenotypic level through motor control, energy, and surprise/cognitive complexity. We propose that these three sets of acoustic features (i.e. control, energy and surprise or CES) combine to form a mathematical calculus of attention that individuals can manipulate to stimulate attentive behavior in an observer (or larger audience). We further propose that the quality of attention signaling can be indicated by the level of non-randomness in the trajectories of sound in the ternary space of CES. We demonstrate that attention signaling defined in this way is common across many forms of human and animal vocalization and also suggest that it is associated with visual displays as well. We investigate auditory attention signaling in human and animal vocalizations compared to other non-biological sounds and demonstrate that stability in CES corresponds to our human perception of musicality in both sounds we culturally define as music, as well as animal vocalizations we perceive as more musical than others (i.e. songbirds, gibbons, and chorus frogs). Overall, we find that professional human musicians have high stability in CES as well as less variance than other animal ‘song’ in their song signatures (i.e. distances derived from functional data analysis of time series trajectories) of control and energy, but not in their signatures of complexity or surprise. This is probably due to the self-selected talent and subsequent training that human singers go through. Unlike chorus frogs, gibbons and songbirds, not all humans have to learn to sing to succeed in life. Our network analysis of song signatures also shows that the genres of opera and jazz are more distinct and well defined (i.e. modular) than popular music, with opera being closer to that of other animals (especially gibbons and chorus frogs, rather than songbirds). This result likely reflects the functional similarity of vocal architecture (i.e. frogs and mammals both have a single larynx while bird use a more complex structured syrinx). Popular music also uses more technological means of achieving near perfect control and energy through amplification, fretboards and keyboards, autotuning, and electronic drums and synthesizers. Jazz relies on less of this and opera even less so. This is also consistent with the observation that we find less individuality of song signature within popular musicians than within other genres (jazz, opera) or within the boundaries of broader taxonomic groups. We also demonstrate that novice vocalists, in both humans and at least one species of bird (Yellow Canary), have lower quality levels of attention signaling than expert individuals as evidenced by increased randomness of song trajectory in the space of CES. This finding is supported by a recent study showing that daily vocal exercise is as important for birds as it is for human singers [1] and therefore is likely to be an honest signal of fitness through performance as well. We also find attention signaling is stronger in musical performances that emphasize the ability of soloists over the perceptions or moods of the listener (i.e. piano concertos vs. ambient piano music). Lastly, we demonstrate that in the career of one famous popular performer (Bjork), live audience feedback seems to constrain the diversity of her exploration of the space of CES when compared to her work in the studio. This suggests there is something consistent to attention signaling that requires audience presence and helps to explain previous observations in neuroscience that show that people are often more stimulated by live vs. recorded music [87].

In summary, we find striking mathematical similarity among animal ‘song’ vocalizations and human musical sounds regarding the stability of CES, along with some well-defined differences across evolutionary distant taxa that evolved singing behavior independently (i.e. anurans, birds, primates) We believe this validates our mathematical definition of fitness in animal vocal performance and human music suggesting further investigations at the intersection of aesthetics and evolutionary biology could prove valuable. In future studies, we plan to investigate the utility of our software and methodology for predicting historic and current trends in the popularity of music (e.g. peak Billboard magazine chart positions, YouTube views, or Spotify downloads) and in training amateur musicians and/or designing new kinds of instrumental or electronic sounds that stimulate the attention of audiences in interesting ways.

Musical sound stimulates the neurophysiology of the human brain more than any other neurological stimulus [75] and may yet hold the answer to some of the yet unanswered questions around the major evolutionary expansion events in hominid brain size[19][28]. We intuitively understand that music affects us greatly in both an external physical sense via motion and attention as well as an internal cognitive sense via emotion and intellectual reflection. The geometry of the calculus of attention provides us with a simple and yet novel way to envision this complex cognitive duality. If we view the ternary plot representing the external acoustic features of musical control, energy and surprise as the foundation for an complementary triangular internal representation of how music might affect our internal awareness, we can build a six-sided star (Figure 10A) that balances (A) the physical impact of music on our motor cortex between the vertices for motor, control and energy, the (B) emotional impact of music between the vertices of energy and surprise (i.e. complexity), and the (C) intellectual or artistic impact of music between the vertices of control and surprise. If we place unbalanced emphasis on the three vertices of control, energy, and surprise, either singularly or in pairs, we can seemingly and yet surprisingly describe the general feelings invoked by hearing different genres of music, simple musical scales, and spoken words (Figure 11). In this internal-external cognitive geometry, the upward triangle might represent the outward focused activity of the ancestral dorsal and ventral attention networks [90] while the downward triangle might represent the inward focus of the more recently evolved default mode network [54].

**Figure 10:**
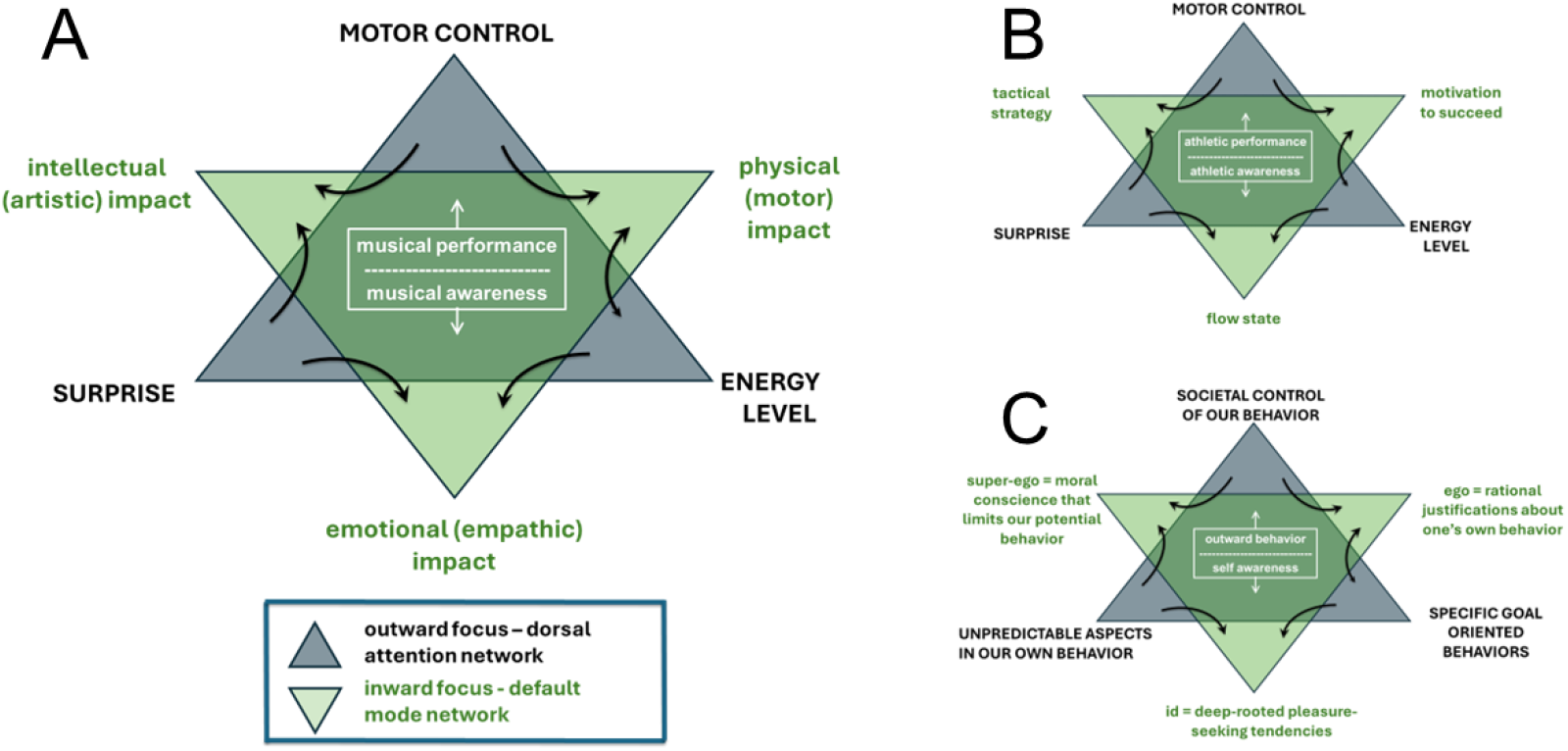
Theoretical relationships between outward and inward attention or focus with respect to (A) music and dance, (B) sports contests, and (C) human social behavior more generally. We theorize that context-dependent features of control, energy, and surprise (gray triangle) act generally to stimulate the dorsal attention network of the brain, while inward attention, likely mediated by the default mode network of the brain, stimulates an inverse geometry of self-awareness (green triangle) specific to that activity. (A) In music and dance, inward self-awareness is balanced between physical, emotional and intellectual (artistic) impacts on the mind. (B) In the context of a sporting event, inward self-awareness is balanced between physical motivation to succeed, emotional flow state and intellectual (tactical) strategy in the mind. (C) In a broader context of all human behavior, features of societal control, individual goal-oriented behavior, and unpredictability in our own and others behavior (i.e. a generalized form of control, energy, and surprise) create an inward geometry of self-awareness that replicates Freud’s concepts of id (i.e. pleasure seeking), ego (i.e. rationalization of one’s own behavior) and super-ego (i.e. moral conscience or judgement of one’s own behavior in the context of societal rules).

**Figure 11:**
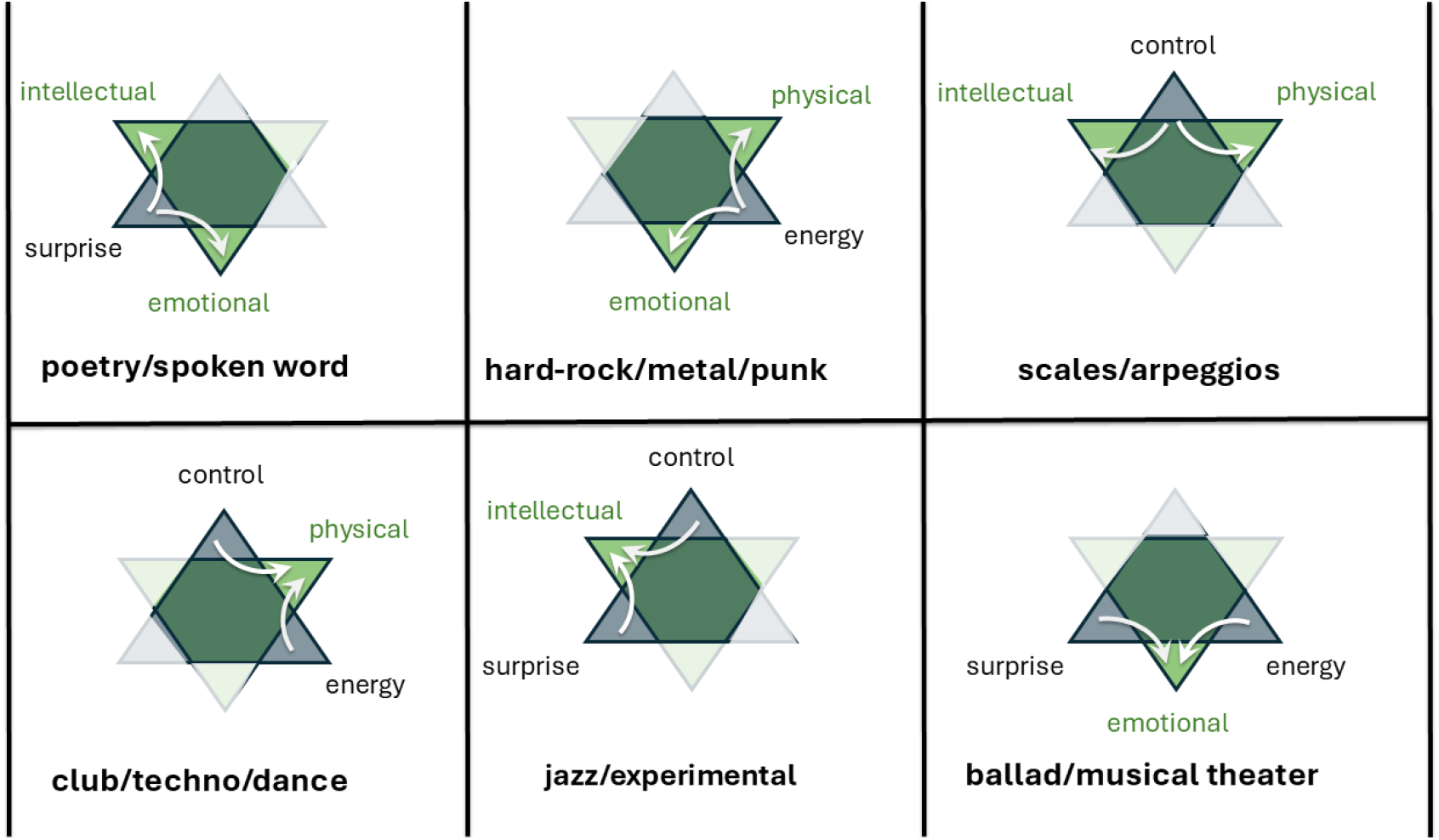
The effects of unbalanced geometry in the calculus of attention (gray) can explain the internal cognitive effects (green) of various musical genres. For example, emphasis on just energy level in music with de-emphasis on novelty/surprise and fine motor control can result in the strong feelings of physical plus emotional impact that we often tend to associate with heavy metal/punk rock. Similarly, a singular emphasis on motor control produces a consequent lack of emotion when we listen to scales or arpeggios, and a singular focus on lyrical novelty, without emphasis energy or motor control explains the cognitive differences we perceive when comparing poetry/spoken word versus music itself. When adjacent pairs of fitness signals are emphasized in combination, it has the effect of focusing on the cognitive impact in a single area. For example, the primarily physical impact we hear in dance music is created by a combination of balanced emphasis on energy level and motor control, with little novelty or surprise. Similarly, the intellectual stimulation of modern jazz is created by the balanced combination of improvisation (surprise) with motor control. And the enhanced emotional impact of song ballads and musical theatre are caused by the balance of energy level and unexpected features in the lyrical or musical elements. Note: not pictured here, popular music would seem to require musicians to maintain balance between all three features of the model (energy, control and surprise).

In the context of music, this geometric model provides a simple demarcation of the difference between outward musical performance and inward musical awareness (Figure 10A). What is also very interesting is that if we consider song to be one of many other possible types of behavioral display, we can apply this geometry much more broadly. For example, like musical performance, any sporting contest observed by an audience might be also be considered an external display of motor control of the body, energy on the field of play, and elements of tactical surprise. These outward observable traits of an athlete relies upon an internal awareness of a physical motivation to succeed, an emotional flow state of body-mind, and an intellectual manipulation of tactics within the rules of the game. (Figure 10B). Most interestingly, this geometry can be broadened even more to include general human social psychology regarding external aspects of societal control of behavior (i.e. control), specific goal-oriented motivations of individuals (i.e. energy), and unpredictability in peoples behavior (i.e. surprise), to reproduce the classic internal Freudian architecture of the id, ego, and super-ego (Figure 10C). While modern neuroscience has largely rejected these Freudian terminologies because of their lack of localization in the brain [57] and studies citing Freudian concepts are in decline [97], interestingly, many clinical psychologists still rely upon them in therapeutic interventions [39] where classic psychoanalysis has evolved into a more modern psychodynamic therapy that is mainly focused upon patient outcome [59]. Our geometric model hypothesizes the existence many potential common features of human and animal behavioral display that manifest from an ancient common calculus of attention in the animal brain. This might underpin our internal self-awareness of not only music, but many other combinations of physical, emotional, and intellectual awareness in response to many other human social behaviors. Perhaps future work within the fields of behavioral ecology and neurophysiology might be able to further confirm or reject the ternary architecture of our mathematical model. Experimental science in general, and neuroscience more specifically, has often struggled with its inability to understand complex emergent phenomena born of simple feedback loops within system dynamics [52]. However, the mathematics of luminaries like Robert May, Ed Lorenz, Benoit Mandelbrot and Mitchell Feigenbaum have demonstrated that complex emergent behavior originating from simple feedback mechanisms are universal in nature, even if not well understood[51][45][21][48]. While our conclusions here might seem tenuous to laboratory scientists trained in reductionist experimental methods based in molecular biology or brain imaging, we would argue that some attempts to understand complex emergent phenomena such as those involved in art and aesthetics can benefit from from some basic theoretical grounding in mathematics when it is combined with data driven computational methods like machine learning. We have attempted to demonstrate this in our work presented here.

In conclusion, we can see network structure between individual songs that strongly reflects the boundaries of taxonomy, culture (i.e. musical genre) and audience presence/absence, but only weakly reflects human individuality within opera, jazz, and popular music when controlling for gender. Thus, our mathematical modeling and comparative analyses of the behavior of human vs. animal ‘song’ are highly suggestive as to how our attentive behaviors and perhaps even some aspects of our self-awareness were probably molded by ancient evolutionary history. The Ediacaran-Cambrian transition 541 million years ago created selective pressures underlying a singular calculus of attention, thereby limiting the dimensionality of our minds in ways that computer algorithms behind machine learning and artificial intelligence are not. Yet, when attention has been implemented in the transformer neural network architecture of modern generative artificial intelligence (GAI), it has vastly improved its capability to mimic and interact with our own neural architecture [88]. The concept of a calculus of attention has broad implications beyond popular music, especially in an emerging internet economy where people’s attention is increasingly monetized for exploitation by GAI [32][89]. We suggest that further investigations of the underlying mathematical patterns and fundamental biology of attentive behavior can potentially help guard our society against GAI applications aimed at overstimulating the cognitive mechanisms that control our attention. Humans not only direct their attention towards popular music, but also popular athletes, public speakers, teachers, preachers, politicians, and newscasters, all of whom might also be exploiting behavioral signals of control, energy, and surprise designed by our shared evolutionary biology to hold our attention. This can potentially affect us in ways that can prove either productive or unproductive to our lives depending upon the context with which we engage in internet outlets for modern media. As the attention and default mode networks of our brains are so strongly anti-correlated in their activity[54], this can perhaps explain why we can too easily lose our sense of self when we are heavily immersed in stimuli that constantly demand our immediate attention.

## Supporting information

examples in mp4

