## Supplemental Figures for "An ancient evolutionary calculus for attention signaling retained in modern music"

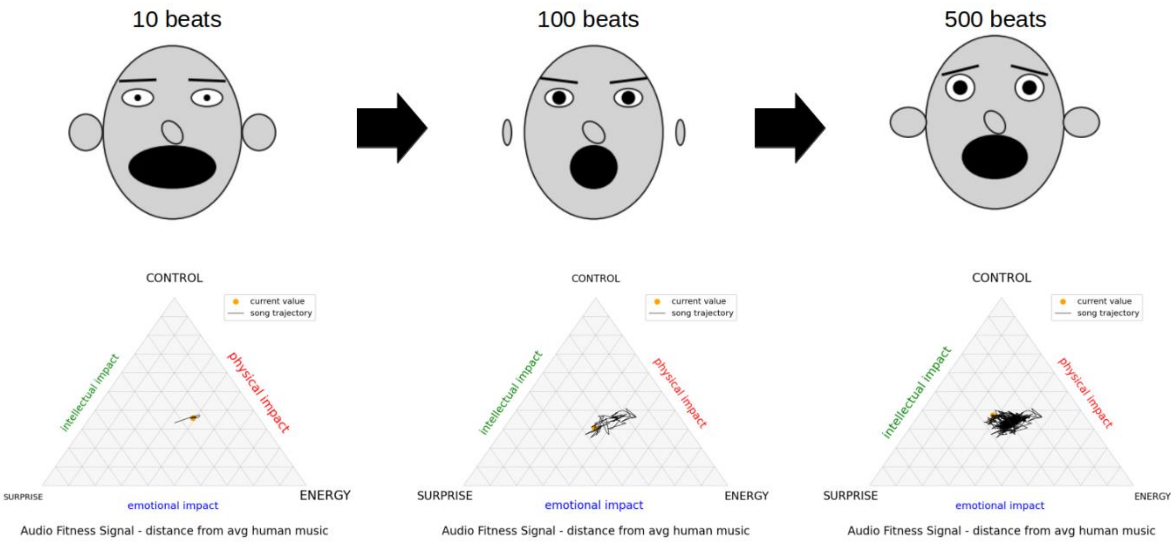


***Figure S1. The POPSTAR software*** *measures three indicators each for sound control (pitch control, timing evenness, and harmonic levels), sound energy (tempo, amplitude modulation, and reverberation), and surprise or sound complexity (note variability index, Lempel-Ziv complexity, multi-scale entropy). It outputs a movie (.mp4) file for each song track as a trajectory within a dynamic ternary or triangle plot in the combined space of control, energy, and surprise (CES) as well as each feature represented by the components of a dynamic Chernoff face, where ear/nose dimensions represent control, mouth/eyebrow dimensions represent energy and eye dimensions represent surprise. The movie is updated according to the user, either on every time beat in the music (determined by beat tracking algorithm) or every ¼ of a second. The fitness signaling in musical sounds are determined through further analysis of the time step distribution of the song trajectory in the ternary plot.*

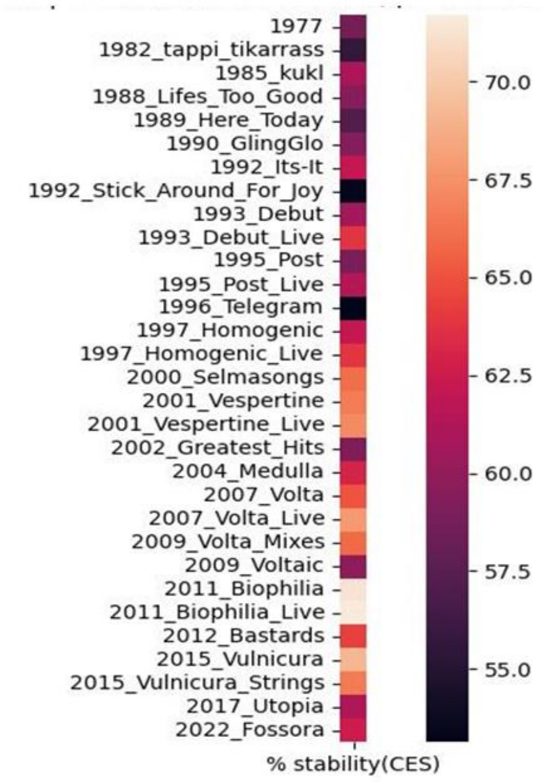


***Figure S2. Stability in control, energy and surprise (CES) across the pop artist Bjork's (Guomundsdottir) entire career from 1977- 2022.***

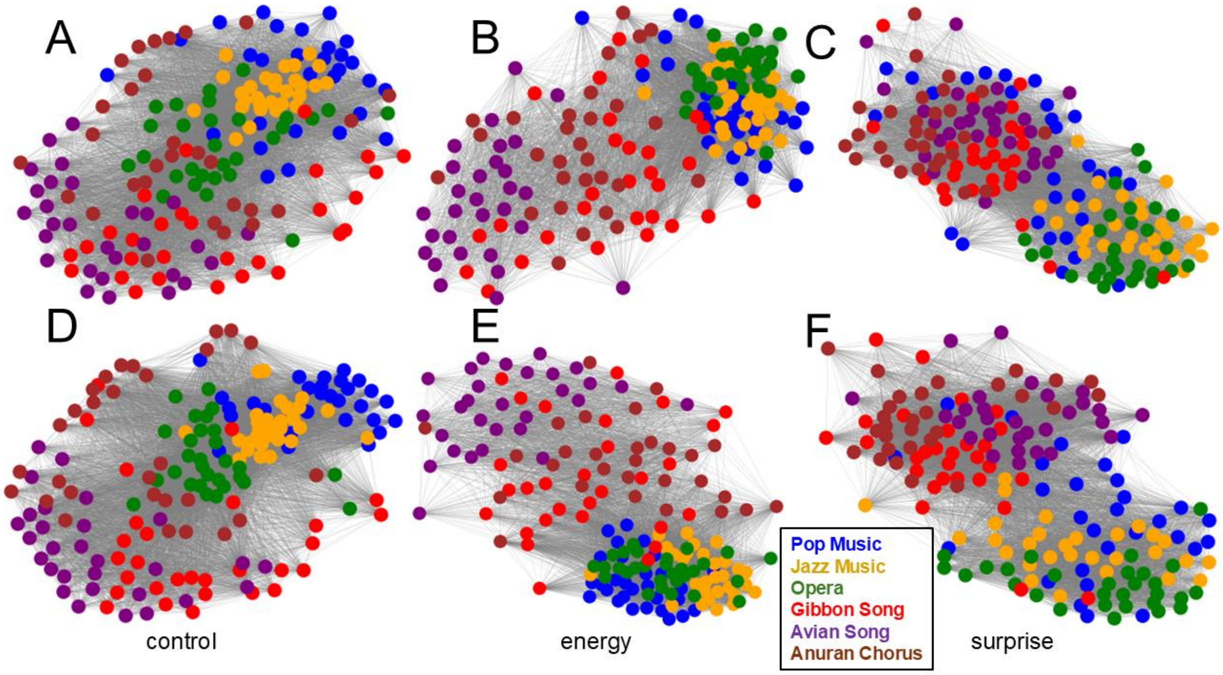


***Figure S3. Adjacency networks of individual songs of frogs, birds, gibbons, and humans mapped within the three dimensions of control, energy and surprise (CES). In A-C the human singers are female and in D-F they are male. In the animals, all singers are assumed to be male except in gibbons (genus Hylobates) where males and females typically sing improvisational duets. Functional data analysis is applied to generate a b-spline interpolation of the time series of an individual song in the specified dimension (either C, E or S). In each network, each node represents an individual song and each edge represents an L2 distance between the functions for two songs. The network representations utilize Kamada-Kawai topology with edges formed using all L2 distances below the median distance observed (i.e. closest 50% of the nodes attract and the farthest 50% of the nodes repulse).***

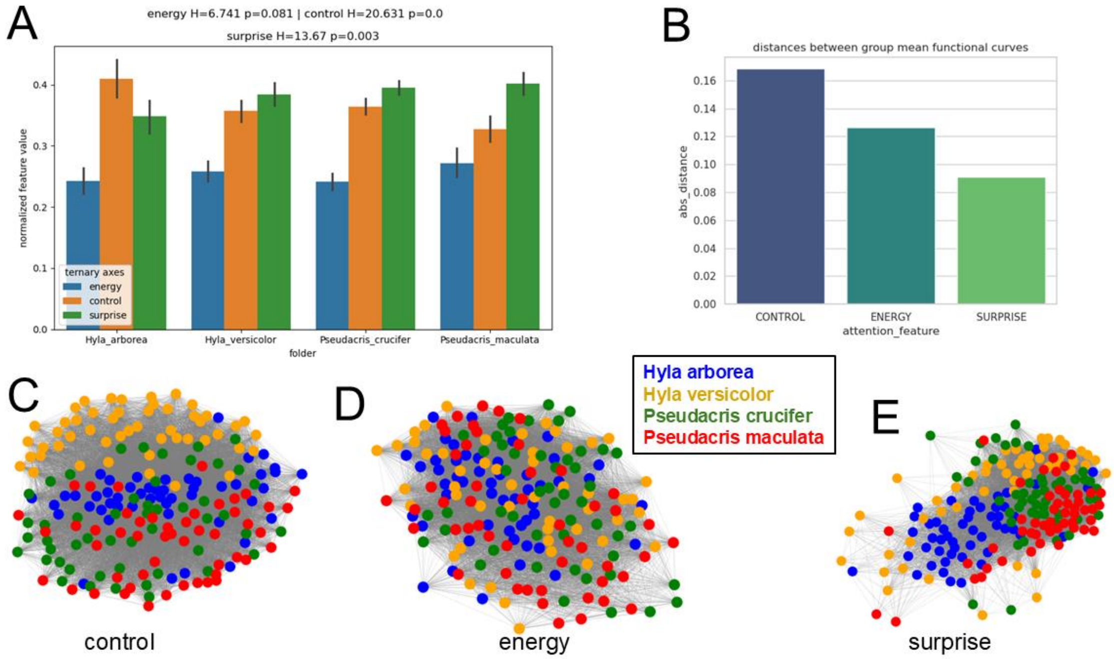


***Figure S4. Across four species of chorus frogs, (A) normalized levels of control, energy and surprise and (B) average L2 distances between individual songs (i.e. variability in song) are shown. Adjacency matrices for individual examples of species songs are also shown for each dimension of (C) control, (D) energy, and (E) surprise.***

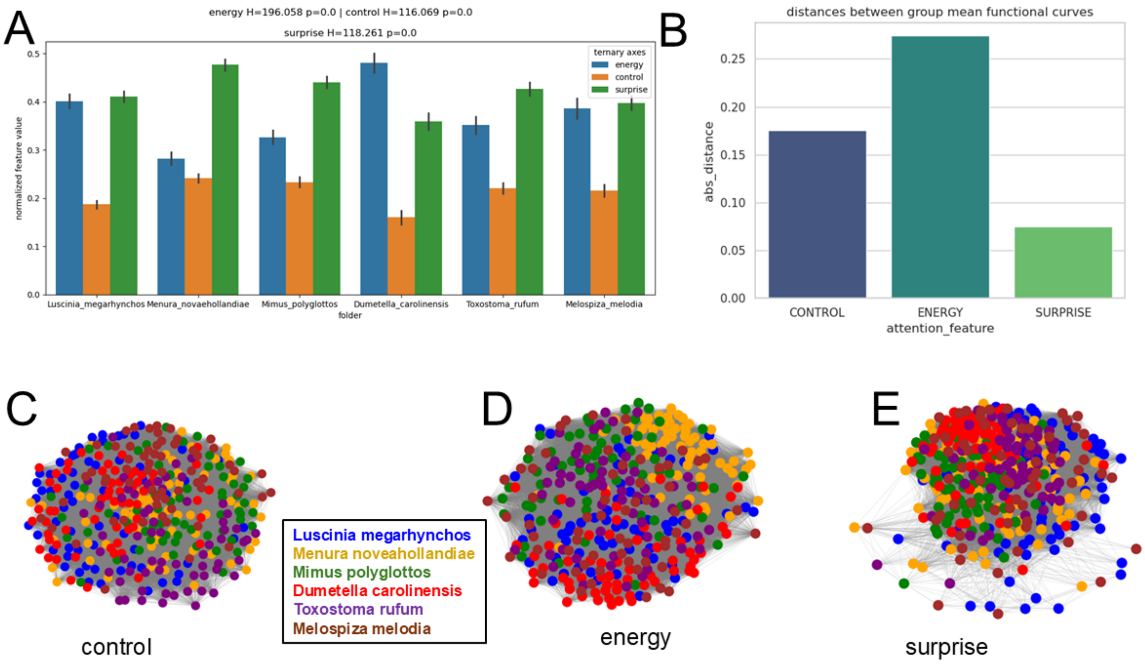


***Figure S5. Across six species of birds with musically complex song, (A) normalized levels of control, energy and surprise and (B) average L2 distances between individual songs (i.e. variability in song) are shown. Adjacency matrices for individual examples of species songs are also shown for each dimension of (C) control, (D) energy, and (E) surprise.***

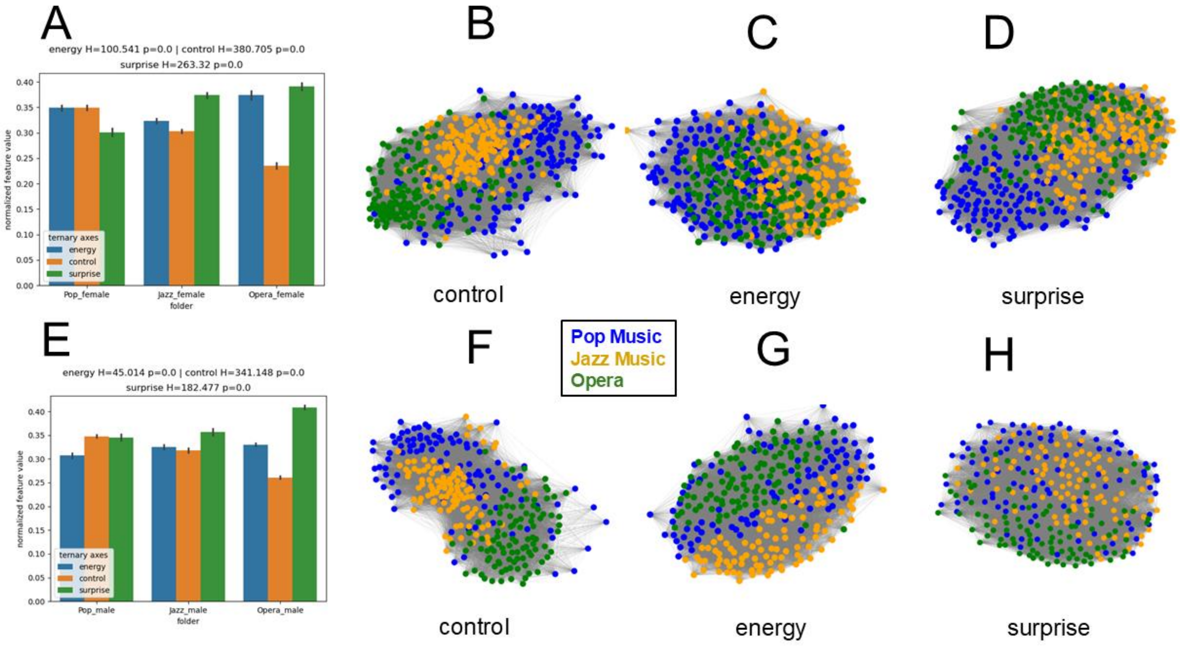


***Figure S6. Across three genres of human music including pop music, jazz  and opera, (A,E) normalized levels of control, energy and surprise and (B-D, F-H) adjacency matrices for individual examples of songs are also shown for each dimension of (B,F) control, (C,G) energy, and (D,H) surprise. Female singers are analyzed in top (A-D and male singers below (E-H).***

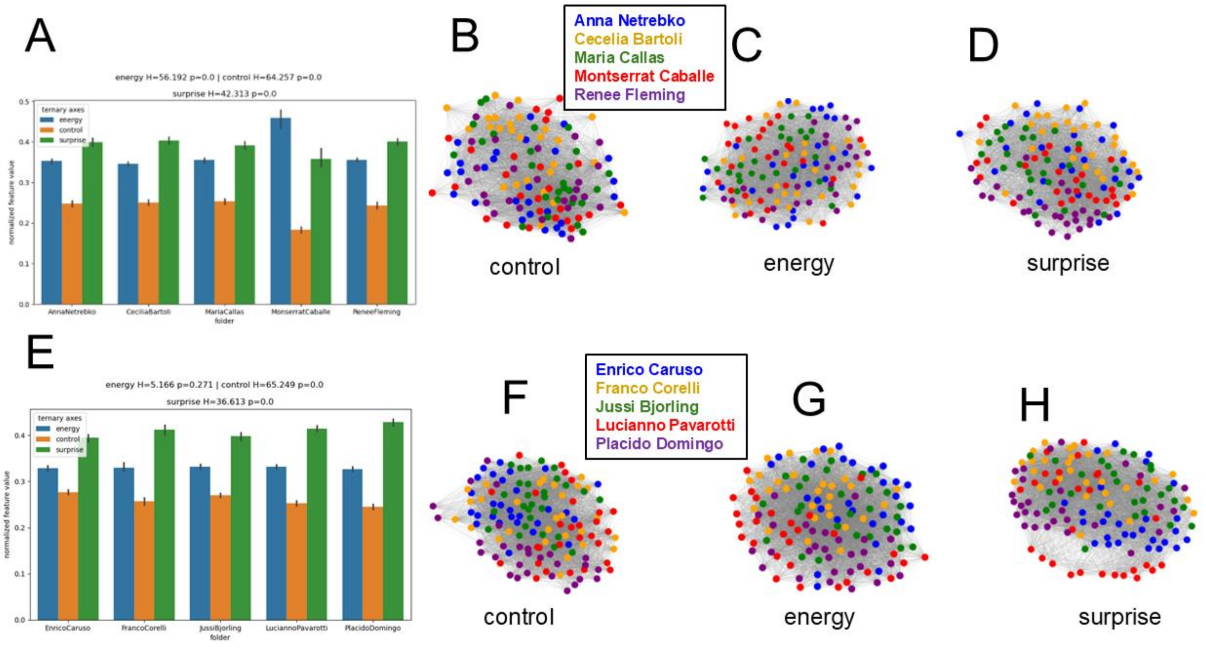


***Figure S7. Across ten famous opera singers, (A, E) normalized levels of control, energy and surprise (B-D, F-H) adjacency matrices for individual examples of opera songs are also shown for each dimension of (B,F) control, (C,G) energy, and (D,H) surprise. Female singers are analyzed in top (A-D and male singers below (E-H).***

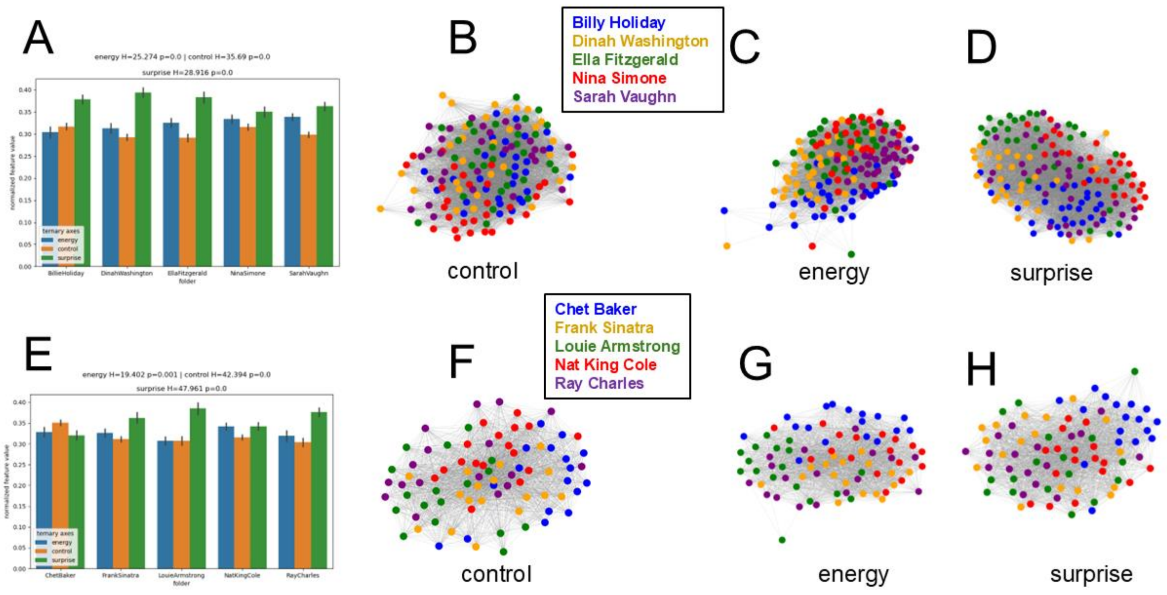


***Figure S8. Across ten famous jazz singers, (A, E) normalized levels of control, energy and surprise (B-D, F-H) adjacency matrices for individual examples of jazz songs are also shown for each dimension of (B,F) control, (C,G) energy, and (D,H) surprise. Female singers are analyzed in top (A-D and male singers below (E-H).***

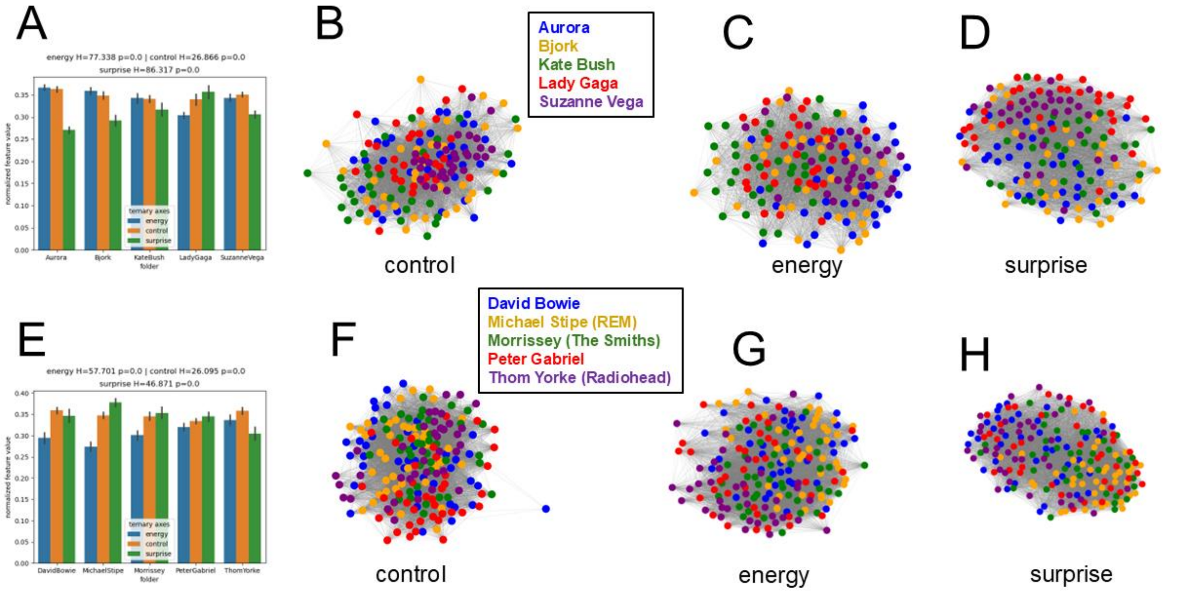


***Figure S9. Across ten well-known pop singers, (A, E) normalized levels of control, energy and surprise (B-D, F-H) adjacency matrices for individual examples of popular songs are also shown for each dimension of (B,F) control, (C,G) energy, and (D,H) surprise. Female singers are analyzed in top (A-D and male singers below (E-H).***

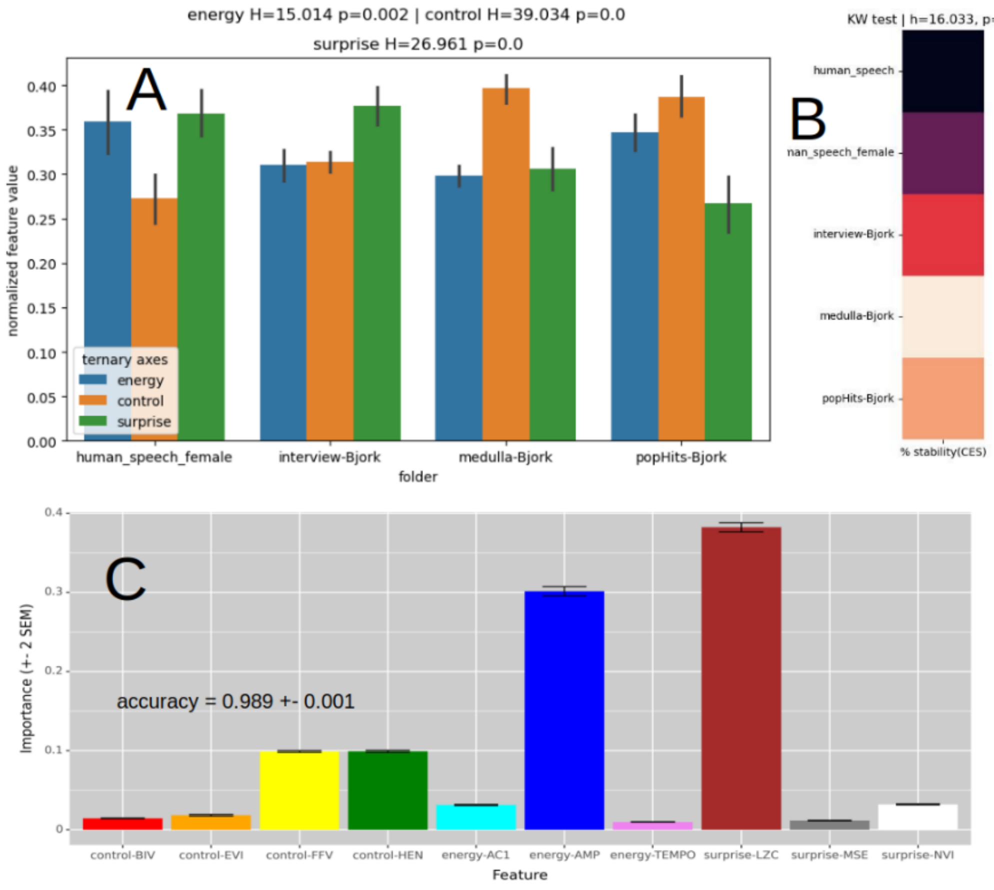


***Figure S10. Comparison of stability in musical control, energy and surprise (CES) in natural speech, music with vocals and instruments, and music created solely with human vocals controlled for a single performer's voice.*** *Here, the Icelandic pop star Bjork is analyzed during her interviews, her greatest hits LP, and her 5th studio album Medulla, which is derived almost entirely with human vocals emulating modern instruments. We see that music has (A) more control and less surprise than speech and (B) more stability in CES than speech, suggesting that the results in Figure 6 could be due to novice singers over-emphasizing patterns normal to speech in their attempts to sing. (C) Random forest classification indicates with high accuracy that differences between Bjork's speech and music are largely driven by Lempel-Ziv (substring) complexity, amplitude modulation, and to a lesser extent, control over pitch and timing.*
